# Inflammatory stress signaling via NF-*k*B alters accessible cholesterol to upregulate SREBP2 transcriptional activity in endothelial cells

**DOI:** 10.1101/2022.05.05.490737

**Authors:** Joseph W. Fowler, Rong Zhang, Bo Tao, Nabil E. Boutagy, William C. Sessa

**Affiliations:** Vascular Biology and Therapeutics Program, Department of Pharmacology, Yale University School of Medicine, 10 Amistad Street, New Haven, CT 06520, USA

## Abstract

There is a growing appreciation that a tight relationship exists between cholesterol homeostasis and immunity in leukocytes, however, this relationship has not been deeply explored in the vascular endothelium. Endothelial cells (ECs) rapidly respond to extrinsic signals, such as tissue damage or microbial infection, by upregulating factors to activate and recruit circulating leukocytes to the site of injury and aberrant activation of ECs leads to inflammatory based diseases, such as multiple sclerosis and atherosclerosis. Here, we studied the role of cholesterol and its master regulator, SREBP2, in the EC responses to inflammatory stress. Treatment of ECs with pro-inflammatory cytokines upregulates SREBP2 cleavage and cholesterol biosynthetic gene expression within the late phase of the acute inflammatory response. Furthermore, SREBP2 activation was dependent on NF-*κ*B DNA binding and canonical SCAP-SREBP2 processing. Mechanistically, inflammatory activation of SREBP was mediated by a reduction in accessible cholesterol, leading to heightened sterol sensing and downstream SREBP2 cleavage. Detailed analysis of NF-*κ*B inducible genes that may impact sterol sensing resulted in the identification of a novel *RELA*-inducible target, *STARD10*, that mediates accessible cholesterol homeostasis in ECs. Thus, this study provides an in-depth characterization of the relationship between cholesterol homeostasis and the acute inflammatory response in EC.

## Introduction

The majority of the biological processes regulating acute inflammation has focused on the contribution of tissue-infiltrating leukocytes. Undoubtedly, leukocytes are crucial for host defense and tissue repair, regulating the balance between resolution and chronic inflammation. However, the endothelium plays a significant role in the overall inflammatory response, particularly in initiation and vascular maintenance. Endothelial cells (ECs) are in constant contact with the bloodstream and rapidly change their phenotype in response to inflammatory stimuli. Inflammatory cytokines, such as tumor necrosis factor alpha (TNF*α*) and interleukin-1 beta (IL1*β*), bind to their respective receptors to activate I-*κ*-kinase, which phosphorylates and degrades inhibitory I*κ*B*α* and releases the key inflammatory transcription factor, NF-*κ*B, to the nucleus (DiDonato *et al.,* 1997). NF-*κ*B, along with other activated transcription factors, such as activator protein 1 (AP1) upregulate the transcription of several inflammatory response genes that increase (1) vascular permeability, (2) leukocyte chemoattraction, and (3) immune cell adhesion and extravasation into tissue (Pober and Sessa, 2007). Indeed, the vascular endothelium is a primary sensor of the circulating bloodstream and is exposed to various stimuli that regulate systemic host defense responses.

It is becoming increasingly appreciated that there exists a connection between cellular immunity and cholesterol. Cellular lipid and cholesterol homeostasis are tightly regulated by the master regulator sterol response element binding factor (SREBP). At sufficient cellular cholesterol levels, SREBP is retained as a full-length protein in the endoplasmic reticulum (ER) bound to adaptor proteins SREBP cleavage-activating protein (SCAP) and inhibitory insulin-induced gene (INSIG) (Brown and Goldstein, 1997). When cellular cholesterol levels decrease, the SCAP-SREBP complex translocates to the Golgi where SREBP is proteolytically cleaved by proteases, S1P and S2P. Cleavage results in the release of the N-terminal fragment of SREBP into the cytoplasm, which translocates to the nucleus to bind to DNA and initiate gene transcription. SREBP1a and SREBP1c isoforms predominantly activate the expression of genes involved in fatty acid synthesis and the SREBP2 isoform upregulates genes that increase cellular cholesterol by de novo synthesis and exogenous uptake (Horton *et al*., 2002).

The relationship between SREBP2, cholesterol homeostasis, and immune phenotype has been predominantly studied in leukocyte immunobiology. First, it has been suggested SREBP2 directly modulates immune responses. In macrophages, it was found that the SCAP/SREBP2 shuttling complex directly interacts with the NLRP3 inflammasome and regulates inflammasome activation via translocation from ER to Golgi (Guo *et al.,* 2018). Another group found that SREBP2 was highly activated in macrophages treated with TNF*α* and that nuclear SREBP2 bound to inflammatory and interferon response target genes to promote an M1-like inflammatory state (Kusnadi *et al*., 2019). Second, several studies have shown that cellular cholesterol levels control immune phenotype. Type I interferon (IFN) signaling in macrophages decreases cholesterol synthesis, allowing for activation of STING on the ER to feed forward and enhance IFN signaling (York *et al.,* 2015). Furthermore, decreasing cholesterol synthesis via *Srebf2* knockout was sufficient to activate the type I IFN response. Mevalonate, an intermediate in the cholesterol biosynthetic pathway, can regulate trained immunity in monocytes. Patients lacking mevalonate kinase accumulate mevalonate and develop hyper immunoglobulin D syndrome (Bekkering *et al.,* 2018). On the other hand, studies have indicated that bacterial lipopolysaccharide (LPS) or type I IFNs can actively suppress synthesis of cholesterol in macrophages and that restoring cholesterol biosynthesis promotes inflammation (Araldi *et al*., 2017; Dang *et al*. 2017).

Despite the importance of the endothelium in the inflammatory response, the link between inflammation and cholesterol homeostasis is not well studied in ECs. There is evidence that increased activation of SREBP2 and cholesterol loading in ECs are pro-inflammatory, but these studies were largely focused on models of atherosclerosis and did not focus on the mechanism of activation, flux of the discrete pools of cholesterol and/or sterol sensing in ECs (Xiao *et al*., 2013; Westerterp *et al*., 2016). Here we show an intimate relationship between inflammatory signaling and cholesterol homeostasis in EC. Tumor necrosis factor *α* (TNF*α*) rapidly activates NF-*κ*B resulting in a time-dependent activation of SREBP2 and SREBP2 dependent gene expression. The activation of SREBP occurs via TNF mediated stimulation of NF-*κ*B and changes in the accessible cholesterol pool that promote sterol sensing but not via other post-translational mechanism. Mechanistically, we show that TNF*α* induction of the NF-*κ*B inducible gene, *STARD10,* in part, mediates the changes in accessible cholesterol leading to heightened SREBP activation. Thus, EC respond to inflammatory cytokine challenges by reducing the accessible cholesterol pool on the plasma membrane thereby inducing canonical SREBP processing and gene expression leading to inflammation.

## Results

### TNFα and RELA transcriptionally regulate canonical SREBP-dependent gene expression

Primary HUVEC were treated with TNF*α* (10ng/mL) for 4 and 10 hr followed by RNA-seq analysis to uncover the transcriptional changes at peak and later stages of activation, respectively. As both a positive control and exploratory aim, we performed RNA sequencing on HUVEC treated with TNF*α* at similar timepoints after RNAi-mediated knockdown of *RELA*, which encodes the protein P65, the key DNA-binding component of the canonical NF*κ*B transcriptional complex. Treatment of HUVEC for 10 hr with TNF*α* resulted in significant upregulation of 913 genes and downregulation of 2202 genes (p < 0.05; −1..5>Fold Change (F.C)>1.5) (Fig. 1a) and *RELA* knockdown decreased expression of 1067 genes and increased expression of 1112 genes (Fig. 1b). As expected, *RELA* gene expression was the most significantly gene decreased after knockdown.

**Figure 1.**
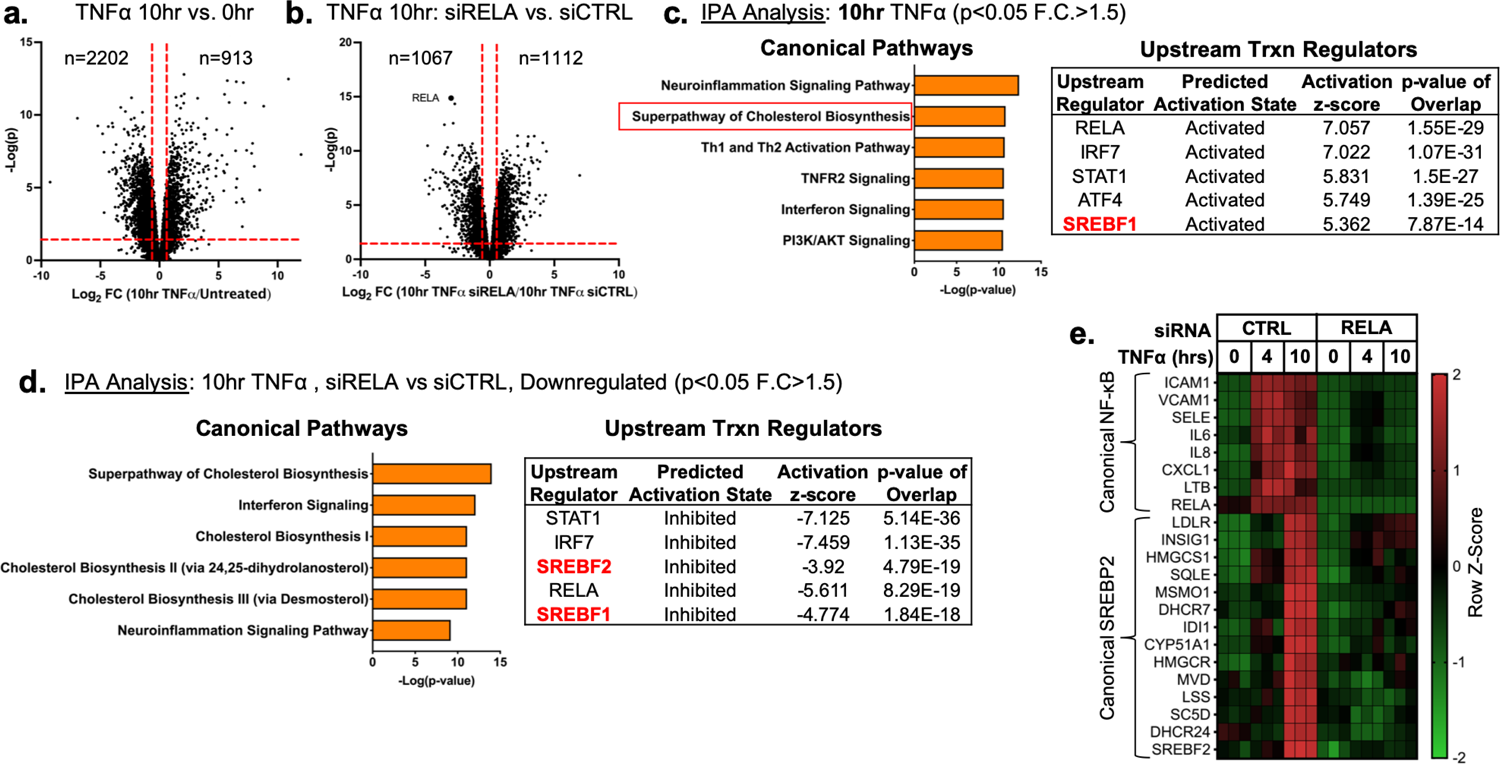
TNF*α* and NF-*κ*B control SREBP2-dependent gene expression in human endothelial cells. Primary HUVEC were treated siRNA against non-targeting sequence (siCTRL) or RELA for 48hr and then incubated with or without 10ng/mL TNF*α* for 10hr. (a) Volcano plot for RNA-seq analysis of differentially expressed genes. Dotted red lines indicate cutoff used for IPA analysis (p<0.05, 1.5<Fold Change (F.C)<-1.5). (b) IPA analysis of most significant canonical pathways and predicted upstream transcriptional regulators for genes that increase at 10hr TNF*α*. (c) IPA analysis of most significant canonical pathways and predicted upstream transcriptional regulators for genes that decrease in cells knocked down with RELA siRNA and treated 10hr TNF*α* compared to control cells treated with 10hr TNF*α*. (d) Representative heatmap of NF-*κ*B and SREBP2 transcriptionally controlled genes from (b) and (c) showing 3 independent donors.

Ingenuity Pathway Analysis (IPA) revealed that TNF*α* treatment for 4hr resulted in the upregulation of several expected pathways reported in literature, including inflammation, TNFR signaling, and activation of IRF (Fig. 1 – Figure Supplement 1 a) (Hogan *et al*. 2017). These pathways were also significantly upregulated in HUVEC treated after 10 hr of TNF*α* treatment (Fig. 1c). Interestingly, Canonical Pathway Analysis uncovered the “Superpathway of Cholesterol Biosynthesis” as the second most significant pathway upregulated in the 10 hr treatment group. Furthermore, Upstream Regulator analysis restricted to transcription factors predicted that SREBF1 was significantly activated in these cells. Metacore analysis of metabolic networks and GSEA hallmark analysis similarly revealed significant upregulation of the cholesterol homeostasis pathway at the 10 hr timepoint (Fig. 1 – Figure Supplement 1 b and c).

RNA-seq pathway analysis of genes reduced after *RELA* knockdown in HUVEC treated with TNF*α* (10hr) revealed that expected inflammatory pathways, such as interferon signaling and neuroinflammation, were significantly inhibited when RELA was not present (Fig. 1d). Additionally, the “Superpathway of Cholesterol Biosynthesis” and several other redundant pathways populated the most significant Canonical Pathway results. Upstream transcription regulator analysis predicted SREBP2 and SREBP1 were significantly decreased in *RELA* knockdown cells. An analysis of gene set overlap between genes significantly upregulated after TNF*α* treatment and genes significantly downregulated in TNF*α*-treated cells lacking *RELA* revealed that SREBP2 target genes were significantly overrepresented (Fig. 1 – Figure Supplement 1 d). We decided to focus on the SREBP2 pathway in late phase (10hr) of TNF*α*-treated cells because of the overwhelming prevalence of cholesterol biosynthesis genes that were increased and significantly attenuated when *RELA* was knocked down (Fig. 1e).

### TNF*α* increases SREBP2 cleavage and transcription of downstream gene targets

RNA-seq analysis predicted that SREBP2 was highly activated in HUVEC treated with TNF*α*. SREBP2 becomes transcriptionally active when its N-terminal DNA-binding fragment is proteolytically processed in the Golgi and allowed to enter the nucleus (Sakai *et al*., 1996). This process can be assayed by measuring precursor (P) and cleaved (C) SREBP2 at 150kDa and 65kDa on a Western Blot, respectively (Hua *et al*., 1995). Measurement of SREBP2(C) throughout a 16 hr timecourse revealed that SREBP2 cleavage began as early as 6 hr after TNF*α* treatment and peaked at 10 hr (Fig. 2a). Furthermore, SREBP2 activation was dose-dependently induced by TNF*α* in HUVEC cultured in either sterol-sufficient fetal bovine serum (FBS) or in lipoprotein depleted serum (LPDS) (Fig. 2 – Figure Supplement 1 a).

**Figure 2.**
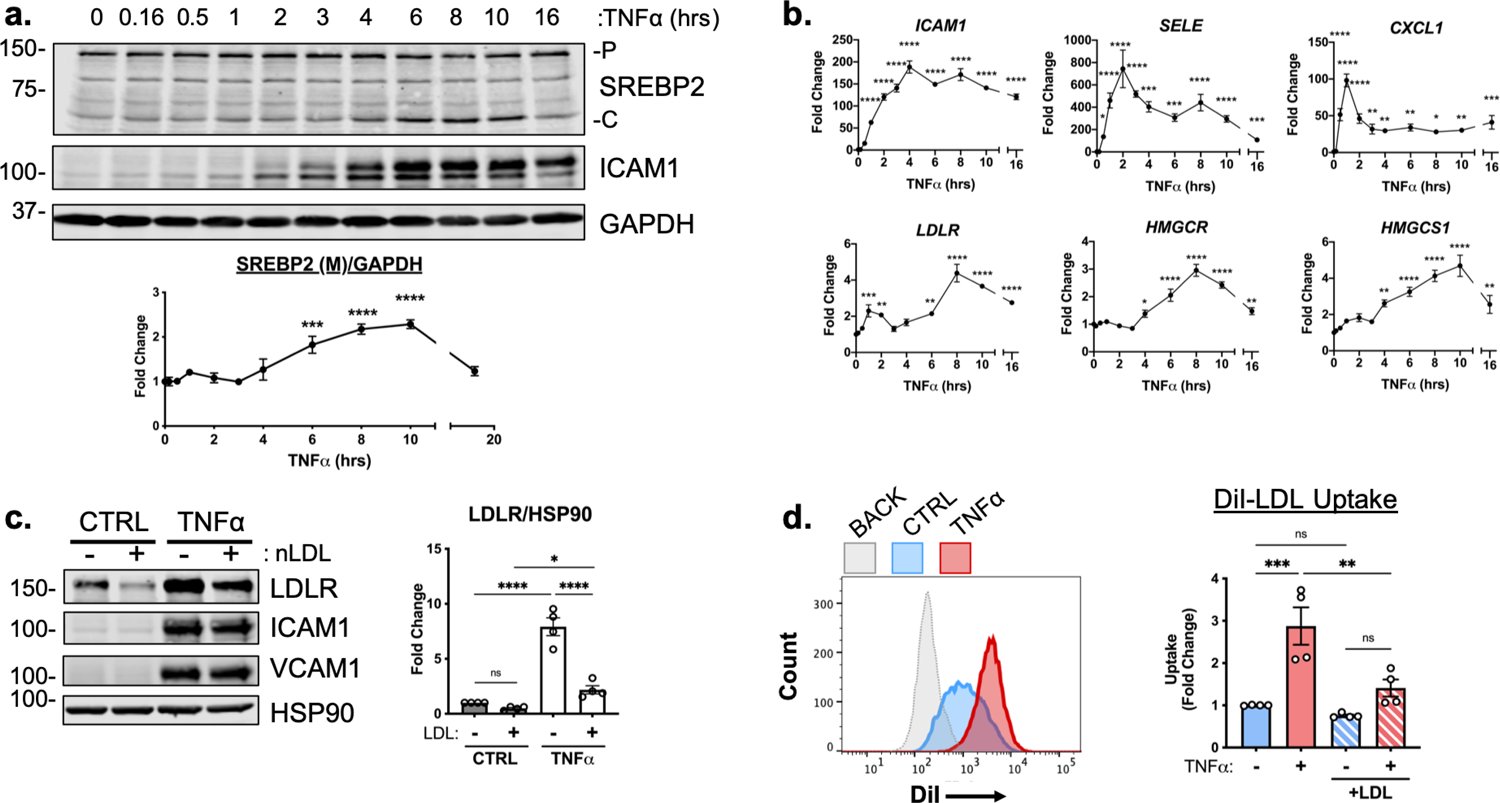
TNF***α*** increases SREBP2 cleavage and transcription of canonical sterol-responsive genes (a) SREBP2 immunoblot from whole-cell lysates from HUVEC treated with TNF*α*(10ng/mL) for indicated time. Data are normalized to respective GAPDH and then to untreated cells (n=3). (b) qRT-PCR analysis of RNA from HUVEC treated with TNF*α* (10ng/mL) for indicated time. Data are normalized to respective *GAPDH* and then to untreated cells (n=8). (c) LDLR protein levels of TNF*α*-treated HUVEC treated with or without native LDL (25μg/mL). Data are normalized to respective HSP90 levels and then to untreated cells (n=4). (d) Flow cytometry analysis of exogenous DiI-LDL uptake in HUVEC treated with TNF*α* and with indicated media. 2.5μg/mL DiI-LDL was incubated for 1hr at 37°C before processing for flow cytometry. Uptake was quantified by PE mean fluorescence intensity per cell and normalized to untreated cells in LPDS across two experiments (10,000 events/replicate, n=4). *p<0.05; **p<0.01; ***p<0.001; ***p<0.0001 by one-way ANOVA with Dunnett’s multiple comparisons test (a and b) or two-way ANOVA with Sidak’s multiple comparisons test (c and d).

We next measured the relative mRNA abundance of NF-*κ*B and SREBP2 target genes throughout the same time course. As expected, known NF-*κ*B target genes, such as *ICAM1, SELE,* and *CXCL1* were rapidly induced after TNF*α* treatment, with many increasing several hundred-fold in less than 2 hr (Fig 2b, top). The patterning of SREBP2 target gene expression was notably several hrs later than NF-*κ*B dependent gene expression. A majority of the canonical SREBP2 genes, including *LDLR, HMGCR, and HMGCS1* significantly increased as early as 6 hr after treatment and peaked at around 8-10 hr (Fig 2b, bottom). Unlike a majority of the cholesterol biosynthesis genes, several fatty acid synthesis genes known to be SREBP1-dependent, such as *ACACA/B, FASN*, and *GPAM* did not significantly increase with TNF*α* treatment and were unaffected by *RELA* knockdown (Fig. 2 – Figure Supplement 1 b).

To further test if SREBP2 activity increased after TNF*α* stimulation, we measured the protein levels of low-density lipoprotein receptor (LDLR), a well-known target of SREBP2 target and receptor involved in the uptake of exogenous lipoproteins, such as low-density lipoprotein (LDL) (Briggs *et al*., 1993). HUVEC were treated overnight in LPDS with or without the addition of 25μg/mL LDL, to suppress SREBP2 cleavage and LDLR expression (Fig. 2c). LDL treatment decreased LDLR protein levels in HUVEC at rest and TNF*α*significantly increased LDLR levels in HUVEC cultured in both LPDS and LPDS+LDL. Exogenous LDL was also able to partially suppress LDLR expression, indicating that this process is sterol-sensitive. Similar results were found for another well-known SREBP2 target of cholesterol biosynthesis, HMGCR (Fig. 2 – Figure Supplement 2 c). Next, we tested fluorescently labeled LDL uptake as a functional readout of the increase in LDLR as quantified by flow cytometry. Similar to what was seen by immunoblotting, TNF*α* treatment led to increased DiI-LDL uptake into cells pre-incubated in various degrees of sterol enriched media (Fig. 2d).

### NF-*κ*B activation and DNA binding are necessary for cytokine-induced SREBP2 cleavage

TNF*α* activates several signaling cascades to fully activate resting endothelial cells that leads to a change in phenotype to promote inflammation. For canonical NF-*κ*B signaling, TNF*α* activates the immediate post-translational activation of the NF-*κ*B complex via phosphorylation and degradation of the inhibitory molecule I*κ*B*α* by I-*κ*-kinase (IKK) isoforms (DiDonato *et al.,* 1997). However, TNF*α* has been shown to upregulate several other signaling pathways, such as JNK, p38, and ERK1/2 (Aggarwal, 2003). Therefore, we sought to confirm that NF-*κ*B signaling is necessary for SREBP2 activation in ECs undergoing inflammatory stress. Treatment of HUVEC with TNF*α*, IL1*β*, and lipopolysaccharide (LPS) to activate NF-*κ*B through separate pathways led to an increase in SREBP2 cleavage and upregulation of LDLR levels (Fig. 3a) with concomitant increases in ICAM1 as a positive control. Furthermore, pre-treatment of HUVEC with the transcription inhibitor, Actinomycin D (ActD), blunted activation of SREBP2, LDLR and ICAM1 in response to TNF*α* (Fig. 3b). This indicated that the mechanism by which inflammatory cytokines activate SREBP2 is most likely through the transcription of novel regulatory molecules rather than via post-translational modifications and/or processing.

**Figure 3.**
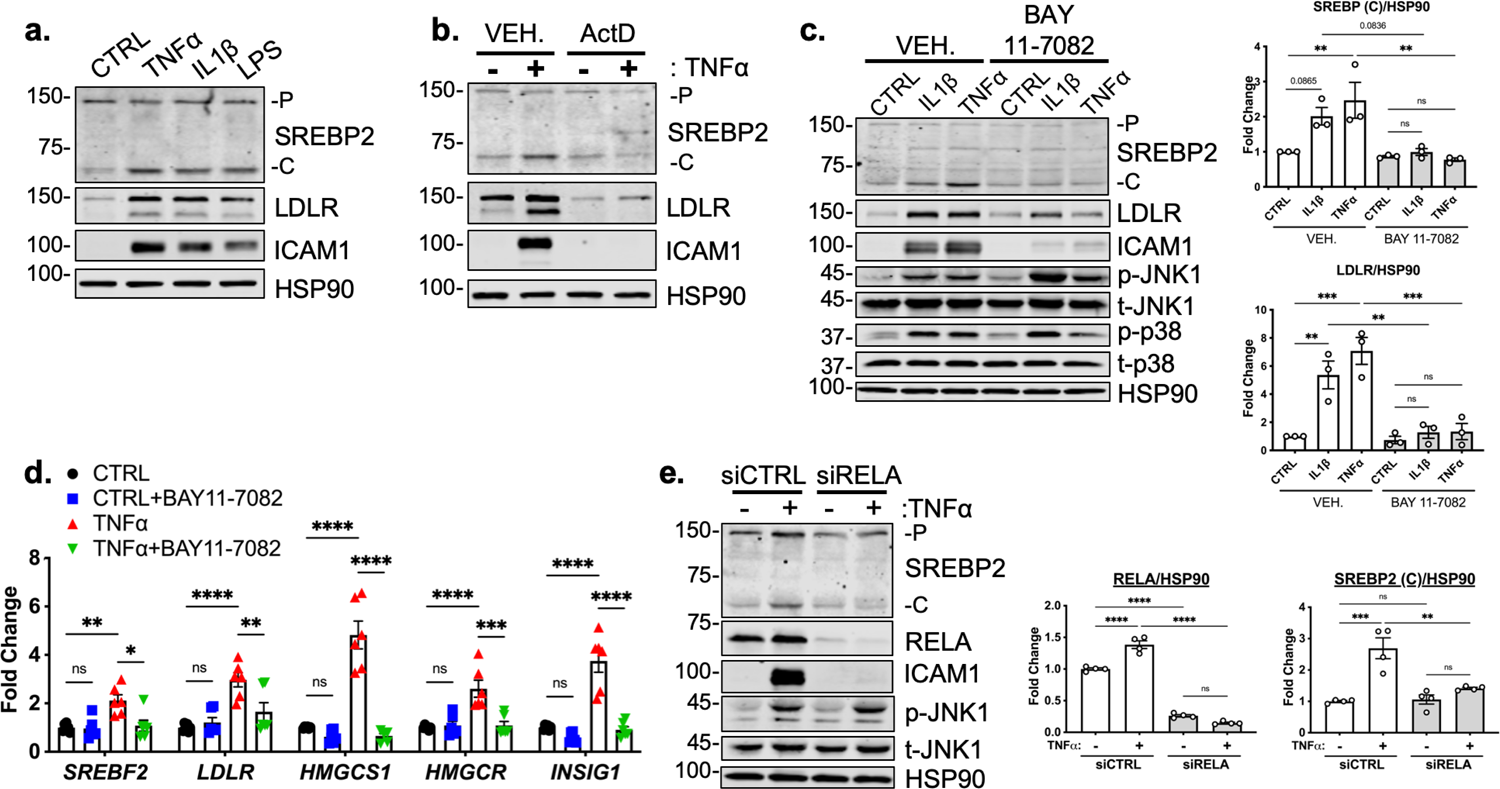
RELA DNA-binding is necessary for activation of SREBP2 by inflammatory stress. (a) Representative immunoblot of SREBP2 and LDLR protein levels in HUVEC treated with TNF*α*(10ng/mL), IL1*β* (10ng/mL), or LPS (100ng/mL). (b) Representative immunoblot of SREBP2 and LDLR protein levels in HUVEC treated with actinomycin D (ActD, 10ng/mL) and with or without TNF*α* (10ng/mL). (c) SREBP2 and LDLR protein levels in HUVEC treated with IL1*β* (10ng/mL) or TNF*α* (10ng/mL) and with or without NF-*κ*B inhibitor, BAY11-7082 (5μM). Data are normalized to respective HSP90 and then to untreated cells (n=3). (d) qRT-PCR analysis of SREBP2-dependent genes, *SREBF2, LDLR, HMGCS1, HMGCR,* and *INSIG1,* expression in HUVEC treated with or without TNF*α*(10ng/mL) and BAY11-7082 (5μM). Data are normalized to respective *GAPDH* and then to untreated cells (n=6). (e) SREBP2 and RELA levels in TNF*α* (10ng/mL)-treated HUVEC treated with or without siRNA targeting RELA. Data are normalized to respective HSP90 and then to untreated cells (n=4). *p<0.05; **p<0.01; ***p<0.001; ***p<0.0001 by one-way ANOVA (c and e) or two-way ANOVA (d) with Tukey’s multiple comparison’s test.

Lastly, we measured TNF*α*-mediated SREBP2 activation after chemical inhibition and genetic knockdown of NF-*κ*B to confirm previous RNA-seq results. Treatment of HUVEC with BAY 11-7082, a selective IKK inhibitor, significantly attenuated IL1*β* and TNF*α* induced increase in SREBP2 cleavage, LDLR protein levels, and mRNA expression of SREBP2-dependent genes (Fig. 3c, 3d) (Keller, *et al*. 2000). Notably, BAY 11-7082 did not suppress JNK or p38 signaling, demonstrating specificity for the NF-*κ*B pathway (Fig 3c). Western Blot analysis of SREBP2 in HUVEC after *RELA* knockdown confirmed that NF-*κ*B DNA-binding and transcriptional activity are required for cytokine induction of SREBP2 cleavage (Fig. 3e).

### Canonical SCAP/SREBP2 shuttling is required for TNF***α***-mediated SREBP2 cleavage

Studies have shown that SREBP2 cleavage can be controlled by mechanisms beyond the SCAP shuttling complex, such as Akt/mTOR/Lipin1 regulation of nuclear SREBP and direct cleavage of SREBP in the ER by S1P (Shimano and Sato, 2017; Kim *et al*., 2018). It is possible that a post-translational SREBP2 regulator could be the RELA-dependent molecule responsible for increased SREBP2 activation. Furthermore, it is also feasible that the *SREBF2* gene itself is under the control of NF-*κ*B, which would upregulate total SREBP2 and increase the threshold of cholesterol needed to suppress its cleavage. Interestingly, this has been reported in a previous study for *SREBF1* (Im *et al.,* 2011). Thus, we used several SREBP-processing inhibitors to test if the SCAP-mediated translocation and Golgi cleavage are necessary for SREBP2 activation in cytokine stimulated EC (Fig. 4a).

**Figure 4.**
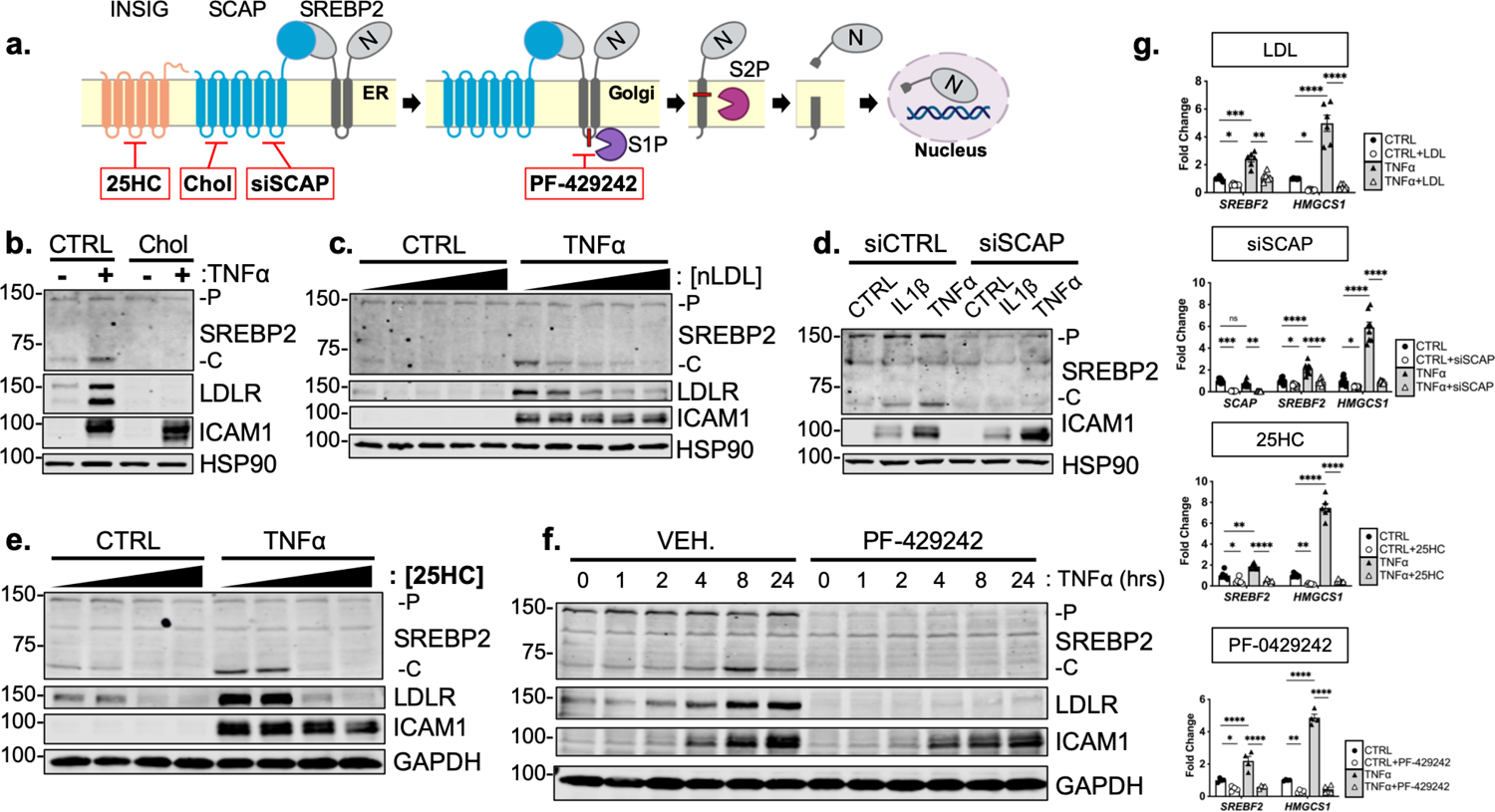
Cytokine-mediated upregulation of SREBP2 cleavage requires proper SCAP shuttling and proteolytic processing in the Golgi (a) Schematic of where 25-hyroxycholesterol (25HC), cholesterol, siSCAP, and PF-429242 inhibit SREBP processing throughout the pathway. (b) Representative immunoblot of SREBP2 and LDLR protein levels in HUVEC treated with TNF*α*(10ng/mL) and cholesterol (Chol) (25μg/mL). Data are normalized to respective HSP90 and then to untreated cells. (c) Representative immunoblot of SREBP2 and LDLR protein levels in HUVEC treated with TNF*α*(10ng/mL) and increasing concentrations of LDL. Data are normalized to respective HSP90 and then to untreated cells. (d) Representative immunoblot SREBP2 cleavage in HUVEC treated with IL1*β* (10ng/mL) or TNF*α*(10ng/mL) and SCAP siRNA. Data are normalized to respective HSP90 and then to untreated cells. (e) Representative immunoblot of SREBP2 and LDLR protein levels in HUVEC treated with TNF*α* (10ng/mL) and increasing concentrations of 25-hydroxycholesterol (25HC). Data are normalized to respective HSP90 and then to untreated cells. (f) Representative immunoblot of SREBP2 and LDLR protein levels in HUVEC treated with TNF*α*(10ng/mL) and PF-429242 (10μM) for indicated time. Data are normalized to respective HSP90 and then to untreated cells. (g) qRT-PCR analysis of *SREBF2, HMGCS1,* and *SCAP* from RNA of HUVECs treated with TNF*α* (10ng/mL) and indicated SREBP2 inhibitor. Data are normalized to respective *GAPDH* and then to untreated cells (n=6). *p<0.05; **p<0.01; ***p<0.001; ***p<0.0001 by two-way ANOVA with Sidak’s multiple comparisons test.

Upon sensing heightened cellular cholesterol, SCAP stabilizes SREBP in the ER and prevents its translocation to the Golgi for processing (Brown and Goldstein, 1997). Therefore, we treated HUVEC with two forms of exogenous cholesterol to test if SCAP shuttling lies upstream of SREBP2 activation in our system: [1] free cholesterol bound to the donor molecule methyl-*β*-cyclodextrin (Chol) and [2] cholesterol-rich LDL. Chol significantly attenuated the increased LDLR and SREBP2 cleavage seen with TNF*α* stimulation (Fig 4b). Furthermore, LDL was able to dose-dependently decrease SREBP2 activation back down to basal levels at the highest concentration of 250μg/mL (Fig. 4c, Fig. 4 – Figure Supplement 1 a). Similar results were seen when SCAP was inhibited with a chemical inhibitor, fatostatin (Fig. 4 – Figure Supplement 1 b). To solidify this point, siRNA knockdown of *SCAP* inhibited the ability of IL1*β* and TNF*α* to upregulate SREBP2 cleavage, but not ICAM1 induction (Fig. 4d).

Next, we sought to inhibit SREBP2 processing by two complimentary approaches, INSIG1-mediated retention in the ER and inhibition of Golgi processing. The oxysterol 25-hydroxycholesterol (25HC) will promote association of INSIG to the SCAP/SREBP2 complex and prevent translocation to the Golgi (Radhakrishnan *et al.,* 2007). Treatment of HUVEC with 25HC significantly prevented cytokine-induced SREBP2 activation and LDLR upregulation (Fig. 4e, Fig. 4 – Figure Supplement c). We next treated the cells with PF-429242, a potent inhibitor of site-1-protease (S1P), which prevented SREBP2 cleavage and LDLR increase throughout the 24 hr timecourse (Fig. 4f, Fig. 4 – Figure Supplement 1 d). The above biochemical experiments were supported by qPCR measurements of SREBP2-dependent genes to confirm that the inhibitors used in this study fully attenuated SREBP2 activity (Fig. 4g). As expected, *HMGCS1* mRNA was depleted basally by LDL, siSCAP, 25HC, and PF-042424 and these compounds prevented the increase in *HMGCS1* transcription in response to TNF*α*. *HMGCS1* transcript levels represented the trend seen in several other sterol responsive genes. Although TNF*α* consistently increased *SREBF2* transcription, all inhibitors also were able to attenuate this upregulation of *SREBF2* mRNA. Taken together, the evidence suggests that canonical SCAP shuttling is necessary for activation of SREBP2 by inflammatory cytokines and that this is not due to direct NF-*κ*B-mediated upregulation of the *SREBF2* transcript or cholesterol biosynthesis genes.

### Inflammatory stress decreases accessible free cholesterol required for SREBP processing

Since SCAP/SREBP2 shutting is maintained when cells were treated with TNF*α*, we reasoned that perhaps TNF*α* regulates cellular cholesterol levels. Lipids were extracted from control and TNF*α*-treated HUVEC and total cholesterol was measured. Incubation of cells in LPDS reduced cellular cholesterol compared to cells cultured in FBS. Cells treated with exogenous methyl-*β*-cyclodextrin-cholesterol (Chol) contained significantly more measured cholesterol (Fig. 5a). However, TNF*α* treatment did not alter total cholesterol in either media condition. Secondly, we quantified cellular cholesterol using mass spectrometry-based lipid analysis, a significantly more precise technique that provides information on molar percentages of lipid. Similar to the initial cholesterol measurements, TNF*α* did not change total cholesterol after 4 or 10 hr of treatment (Fig. 5b).

**Figure 5.**
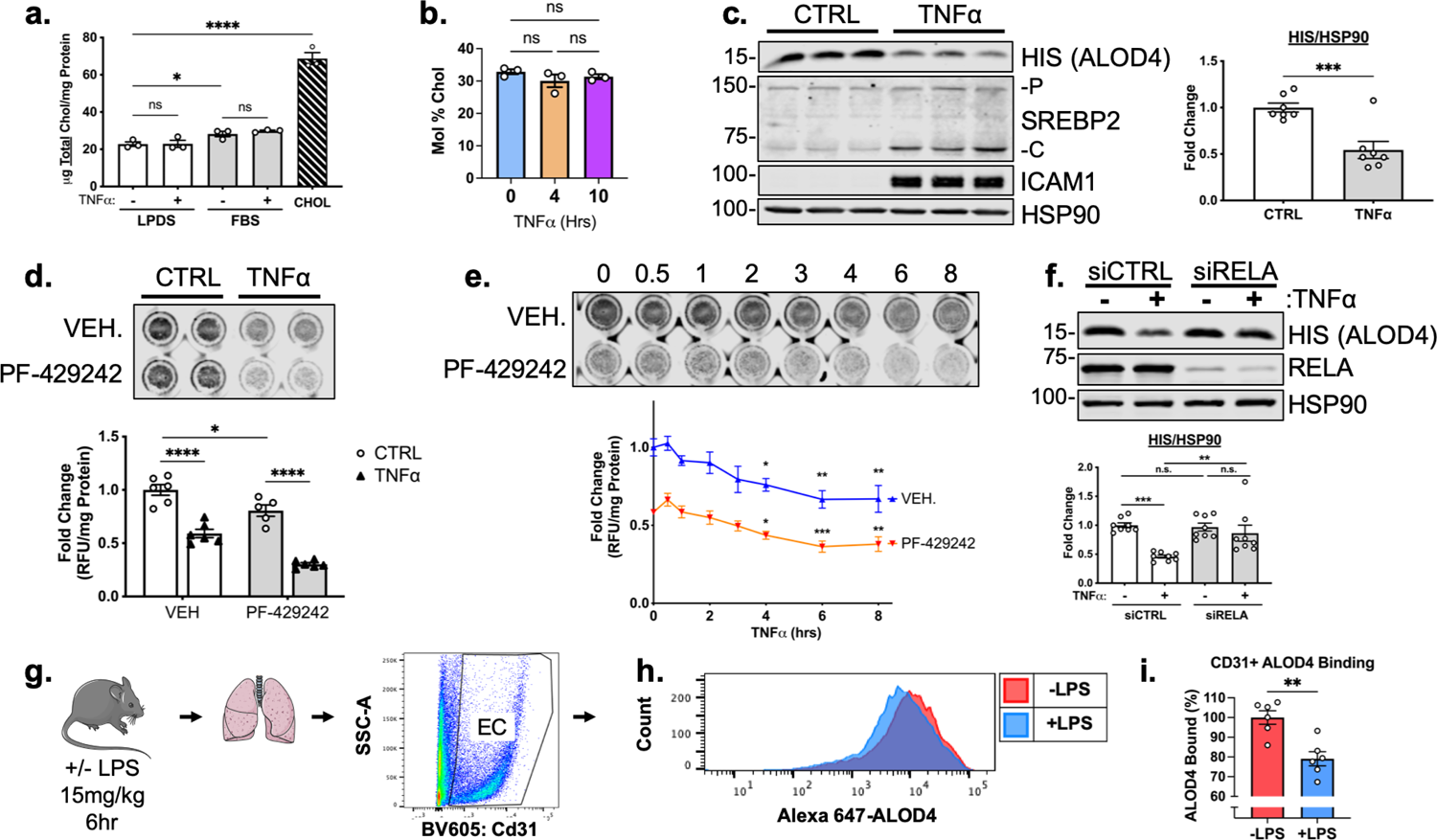
TNF*α* decreases accessible cholesterol in cultured HUVEC and mouse lung ECs *in vivo* (a) Quantification of total cholesterol extracted from HUVEC treated with or without TNF*α*(10ng/mL) and indicated positive controls, lipoprotein deficient serum (LPDS), fetal bovine serum (FBS), or M*β*CD-cholesterol. Data were normalized to respective total protein (n=3). (b) Total cholesterol in HUVEC after 4 or 10 hr of TNF*α* (10ng/mL) quantified by mass spectrometry (n=3). (c) ALOD4 protein levels in HUVEC treated with TNF*α* (10ng/mL). Data are normalized to respective HSP90 and then to untreated cells (n=7). (d) In-cell Western blot of ALOD4 protein levels in HUVEC treated with TNF*α* (10ng/mL) and PF-429242 (10μM). Data are normalized to respective total protein and then to untreated cells (n=6). (e) In-cell Western blot of ALOD4 protein levels in HUVEC treated with TNF*α* (10ng/mL) and PF-429242 (10μM) for indicated time. Data are normalized to respective total protein and then to untreated cells (n=6). (f) ALOD4 protein levels in TNF*α* (10ng/mL)-treated HUVEC treated with or without RELA siRNA. Data are normalized to respective HSP90 and then to untreated cells (n=8). (g) Schematic of protocol to isolate mouse lung endothelial cells and quantify ALOD4 binding by flow cytometry. (h) Representative histogram of ALOD4 binding in Cd31+ lung endothelial cells in mice treated with or without LPS (15mg/kg) for 6 hr. (i) Quantification of ALOD4 binding across 2 flow cytometry experiments in mice treated with or without LPS (15mg/kg). Binding was quantified as AlexaFluor647 mean fluorescent intensity per cell (100,000 events/replicate). Data are normalized to nontreated mice (-LPS, n=6; +LPS, n=6). *p<0.05; **p<0.01; ***p<0.001; ***p<0.0001 by one-way ANOVA with Tukey’s multiple comparison’s test (a and d) or Dunnett’s multiple comparisons test (e), unpaired t-test (c and i), or two-way ANOVA with Sidak’s multiple comparisons test (f).

Changes in the distribution of cholesterol could account for SREBP2 activation without loss in total cholesterol mass. Recently, several tools have been developed to analyze the exchangeable pool of free cholesterol that exists in flux between the ER and the plasma membrane and this accessible pool tightly regulates the shuttling of SCAP/SREBP2 (Infante and Radhakrishnan, 2017). Accessible cholesterol can be quantified using modified recombinant bacterial toxins that bind in a 1:1 molar ratio to accessible free cholesterol on the plasma membrane (Gay *et al*. 2015). To examine accessible cholesterol in EC, we purified His-tagged anthrolysin O (ALOD4) and used it as a probe (Endapally *et al*. 2019). To assess the utility of the probe in HUVEC, cells were incubated with Chol, LDL or with methyl-*β*-cyclodextrin (M*β*CD, a cholesterol acceptor) and ALOD4 bound was quantified by probing for anti-HIS at 15kDa by Western blotting. As expected, treatment with Chol or LDL increased ALOD4 binding, whereas M*β*CD decreased ALOD4 binding (Fig. 4 – Figure Supplement 1 a). Secondly, a similar method was used, but instead of cell lysis, DyLight680-conjugated anti-HIS antibody was directly applied to the ALOD4-incubated cells and read live on LICOR Biosciences Odyssey CLx platform (In-Cell Western Blot). Likewise, the positive controls were able to tightly regulate ALOD4 binding and fluorescence signal (Fig. 4 – Figure Supplement 1 b).

TNF*α* treatment of HUVEC significantly decreased ALOD4 binding (Fig. 5c). Using In-Cell Western blotting, treatment with PF-429242 to reduce SREBP2 processing and its transcription decreased ALOD4 basally and, when combined with TNF*α*, significantly decreased accessible cholesterol even further (Fig. 5d). This suggests that the decrease in accessible cholesterol was independent of SREBP2 stability. Probing accessible cholesterol throughout an 8 hr timecourse revealed that ALOD4 binding significantly decreased as early as 4 hr after TNF*α* treatment in HUVEC treated with and without PF-429242 (Fig. 5e). This was in line with previous results measuring SREBP2 cleavage and gene expression as accessible cholesterol depletion should precede SREBP2 activation. Moreover, *RELA* knockdown attenuated the decrease in accessible cholesterol, indicating that SREBP2 activation by NF-*κ*B was most likely through upregulation of a molecule or pathway that decreases accessible cholesterol (Fig. 5f).

Lastly, we validated that inflammatory stress decreased accessible cholesterol not only in cultured HUVEC, but also in EC *in vivo.* We directly labeled ALOD4 with AlexaFluor 647 and incubated this probe with suspended HUVEC to validate the flow cytometry assay (Fig. 4 – Figure Supplement 1 c and d). Modulation of cholesterol with various treatments altered ALOD4 binding as expected and treatment of HUVEC with TNF*α* revealed similar results to immunoblotting (Fig. 4 – Figure Supplement 1 e and f). Next, we intraperitoneally injected wildtype C57BL/6J mice with a nonlethal dose of LPS at 15mg/kg to stimulate a systemic inflammatory response. Indeed, TNF*α* peaked in the serum of these animals 2 hr after injection (Fig. 4 – Figure Supplement 1 g). Lungs were harvested 6 hr after LPS injection and cells were broken up into a single-cell suspension for flow cytometry staining (Fig. 5g). Cd31+ ECs from mice treated with LPS contained about 20% less accessible cholesterol compared to ECs from untreated mice (Fig. 5h, 5i). Notably, total serum cholesterol remained unchanged in TNF*α*-treated animals compared to control, indicating that the decrease in accessible cholesterol reflects the effect of the inflammatory stimulus on ECs (Fig. 4 – Figure Supplement 1 h).

### STARD10 is necessary for full SREBP2 activation by TNFα

TNF*α* did not impact several canonical biological pathways that regulate the pool of accessible cholesterol. Briefly, cholesterol efflux, sphingomyelin shielding, esterification, and lysosomal/endosomal accumulation remained unchanged in ECs treated with TNF*α* (Fig. 5 – Figure Supplement 1). Therefore, we probed our RNA-seq dataset for genes that have been reported to regulate lipid dynamics and could possibly be upstream of the depletion in accessible cholesterol.

We specifically isolated genes that significantly increased after 4 or 10 hr TNF*α* treatment and decreased with *RELA* knockdown. We found several genes that perform various lipid-associated functions, such as direct lipid binding and transport (*STARD4, STARD10,* and *ABCG1*), free fatty acid enzymatic activation and transport (*ACSL3, DBI*), mediation of mitochondrial steroidogenesis (*DBI, ACBD3, NCEH1*), and metabolism of phospholipids (*SGPP2, PAPP2A,* and *PPAPP2B*) (Fig. 6a). Furthermore, several of these genes have been previously reported to have RELA bound to their respective promoters in ECs treated with IL1*β* or TNF*α*, which supported the hypothesis that these genes were targets for NF-*κ*B (Hogan *et al*., 2017).

**Figure 6.**
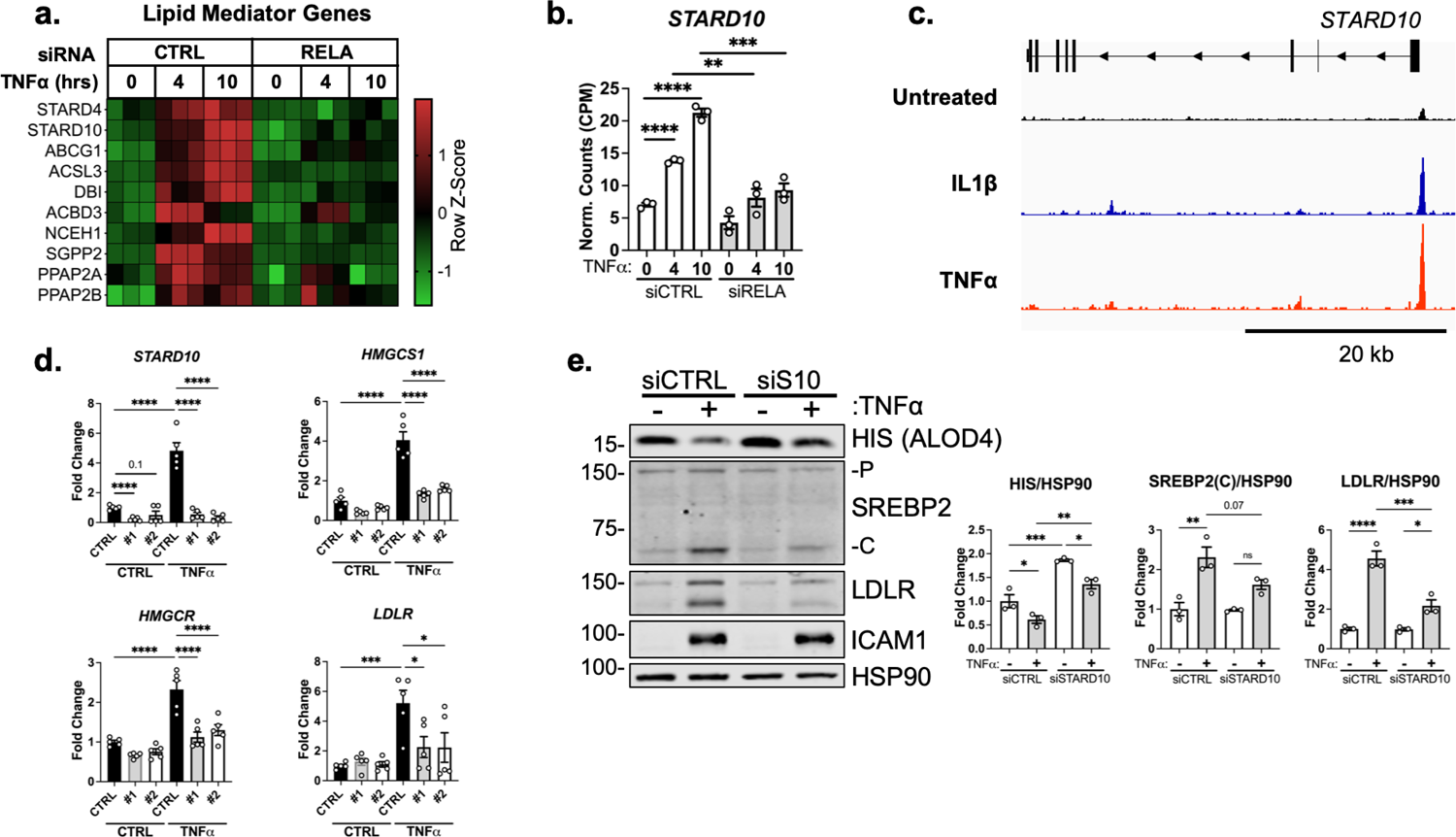
STARD10 is necessary for complete TNF*α*-mediated accessible cholesterol reduction and SREBP2 activation (a) Heatmap of genes that regulate lipid homeostasis, significantly increased with TNF*α* (10ng/mL) treatment after 4 or 10 hr, and were significantly inhibited by RELA knockdown. (b) Normalized counts of *STARD10* expression from previous RNA-seq experiment. (c) *STARD10* gene locus from P65 ChIP-seq analysis of human aortic endothelial cells (HAEC) treated with TNF*α* (10ng/mL) or IL1*β* (2ng/mL) for 4 hr. Data originated from GSE89970. (d) qRT-PCR analysis of RNA from HUVEC treated with TNF*α*(10ng/mL) and two independent siRNA targeting *STARD10* (#1, #2). Data are normalized to respective *GAPDH* and then to untreated cells (n=5). (e) Immunoblot of ALOD4, SREBP2, and LDLR protein levels in HUVEC treated with STARD10 siRNA (siS10) and with or without TNF*α* (10ng/mL). Data are normalized to respective HSP90 levels and then to untreated cells (n=3). *p<0.05; **p<0.01; ***p<0.001; ***p<0.0001 by two-way ANOVA with Sidak’s multiple comparisons test (d and e)

From these several targets identified by RNAseq, we identified *STARD10* as a promising upstream mediator of accessible cholesterol in ECs treated with inflammatory cytokines. STARD10 belongs to a family of proteins that bind hydrophobic lipids via a structurally conserved steroidogenic acute regulatory-related lipid transfer (START) domain (Clark, 2020). STARD proteins regulate non-vesicular trafficking of cholesterol, phospholipids, and sphingolipids between membranes. *STARD10* was found to be upregulated by TNF*α* after 4 and 10 hr of treatment and inhibited by loss of *RELA* (Fig. 6b)*. STARD4* also shared this expression pattern, however, it is a sterol-sensitive gene and its upregulation by TNF*α* most likely occurred via SREBP2 activation (Breslow *et al*., 2005). Analysis of publicly available RELA chromatin immunoprecipitation (ChIP) sequencing data revealed that the *STARD10* promoter contained a strong RELA binding peak in ECs treated with IL1*β* or TNF*α*(Fig. 6c). STARD10 has been shown to bind phosphatidylcholine (PC), phosphatidylethanolamine (PE), and phosphatidylinositol (PI), but much of its detailed biology remains unknown (Olayioye *et al.,* 2005; Carrat *et al*. 2020). Although it has not been shown to directly bind cholesterol, STARD10 may be implicated in the reorganization of membrane phospholipids that could alter cholesterol flux and shield cholesterol localization (Tabas, 2002; Mesmin and Maxfield, 2009; Lagace, 2015).

We next knocked down STARD10 to analyze its role in cholesterol homeostasis and EC inflammatory response. Treatment of ECs with two independent siRNAs targeting STARD10 significantly decreased *STARD10* expression and attenuated the enhanced expression of SREBP2 target genes *HMGCS1, HMGCR,* and *LDLR* (Fig. 6d). Furthermore, STARD10 knockdown significantly rescued the loss in accessible cholesterol that occurs with TNF*α* stimulation (Fig. 6e). SREBP2 activation and LDLR upregulation were also attenuated with STARD10 silencing. Therefore, we have identified a novel RELA-inducible gene in ECs that mediates the cholesterol homeostasis in response to inflammatory stress.

## Discussion

Little is known about cholesterol metabolism and homeostasis in ECs in the context of physiology or pathology. Here, we show that TNF*α* and IL-1*β* induce a transcriptional response via NF-*κ*B that alters cholesterol homeostasis in ECs (Fig. 7). Accordingly, rapid changes in accessible cholesterol then activates SREBP2 processing to compensate for the altered flux of accessible cholesterol between the plasma membrane and ER. Importantly, we identified a novel NF-*κ*B inducible gene, *STARD10*, that serves as an intermediate bridging TNF activation to changes in accessible cholesterol and SREBP2 activation. Collectively, these data support the growing evidence for an intimate relationship between inflammatory signaling and cholesterol homeostasis.

**Figure 7.**
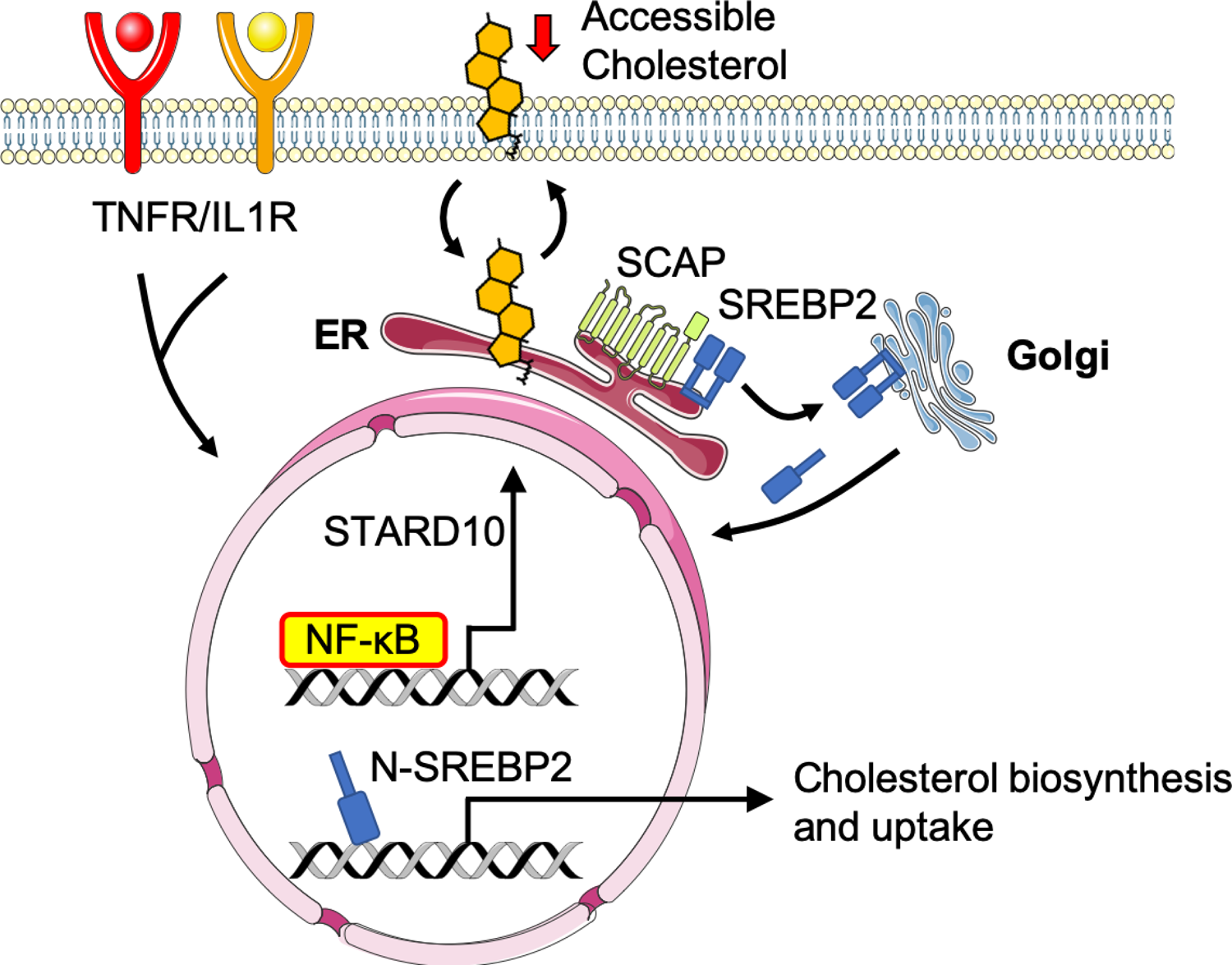
Working model of the relationship between sterol sensing and EC acute inflammatory response Pro-inflammatory cytokines, such as TNF*α* and IL1*β*, promote NF-*κ*B activation of gene transcription in endothelial cells. NF-*κ*B upregulates factors, such as *STARD10*, that significantly decrease accessible cholesterol on the plasma membrane. SCAP senses the reduction in accessible cholesterol and shuttles SREBP2 to the Golgi to initiate classical proteolytic processing. Active N-SREBP2 translocates to the nucleus to transcriptionally upregulate canonical cholesterol biosynthetic genes.

Different forms of cellular stress have been shown to activate SREBP2 in multiple cell types, including oscillatory shear stress in ECs, ER stress in hepatocytes, and TNF*α* in macrophages (Xiao *et al*., 2013; Kim *et al*., 2018; Kusnadi *et al.,* 2019). However, these studies did not implicate changes in cholesterol accessibility as an upstream mechanism leading to the activation of SREBP2. Here, we clearly demonstrate that activation of NF-kB leads to changes in the accessible cholesterol pool promoting classical SREBP2 processing in a SCAP-dependent manner. Epithelial cells treated with IFN*γ*-stimulated macrophage conditioned media rapidly reduce accessible cholesterol to protect against bacterial infection by preventing cholesterol-mediated transport between cells (Abrams *et al*., 2020). Similarly, macrophages deplete accessible cholesterol to protect against bacterial toxin-mediated injury (Zhou *et al*., 2020). In both of these studies, accessible cholesterol was mediated by IFN-induction of the enzyme, cholesterol 25-hydroxylase (Ch25H), which produces 25-HC as a product and enhances the esterification of cholesterol. However, Ch25H is not expressed ECs with or without TNF*α* and cholesterol does not mobilize into the lipid droplet pool. Nonetheless, EC depletion of accessible cholesterol may be an evolutionary mechanism of host immunity.

In the present study, we show that TNF*α* activation of NF-*κ*B, induces the expression of the gene, STARD10. Although little is known about the physiological function of STARD10, it may influence cholesterol homeostasis in several ways. Firstly, it has been reported that STARD10 can directly bind PC, PE, and PI and may alter intracellular membrane dynamics to influence the flux of cholesterol. It has been suggested PC plays a role in cholesterol sequestration either by steric hindrance from by its polar head group or through the creation of novel membranes that may sink cholesterol out of the accessible pool (Mesmin and Maxfield, 2009; Lagace, 2015). Furthermore, STARD10 belongs to a classical family of lipid transporters and may bind lipids not previously reported. As a novel NF*κ*B inducible gene, STARD10 regulation of cholesterol homeostasis is a novel concept that warrants further investigation.

Previous studies indicate that SREBP2 activation feeds forward into the inflammatory response and exacerbates inflammatory damage. Firstly, SREBP2 has been shown to regulate inflammatory phenotype via modulation of cholesterol homeostasis. Increased cholesterol flux has been reported to feed into multiple immune pathways, such as interferon responses, inflammasome activation, and trained immunity (York *et al*., 2015; Dang *et al*., 2017; Bekkering *et al.,* 2018). Furthermore, perturbations in cellular cholesterol may change membrane dynamics and affect cellular signaling (Araldi *et al*., 2017). Secondly, SREBP2 has been proposed to bind and promote transcription of several pro-inflammatory mediators, such as *IL1β*, *IL8, NLRP3,* and *NOX2 (*Kusnadi *et al*., 2018; Xiao *et al.,* 2013; Yeh *et al.,* 2004). SREBP2 binding to non-classical gene promotors may very well depend on cellular and epigenetic context. How SREBP2 feeds into post-translational inflammatory response phenotype of ECs, such as permeability, sensitivity to infection, and receptor signaling, remains to be explored.

This study extends a growing body of work identifying a tight relationship between cholesterol homeostasis, inflammation and immunity. Although we have shown EC response to inflammatory cytokines in the acute setting, much remains to be explored in the context of chronic inflammation. Of particular interest, it has been well appreciated that the endothelium plays an important role in the progression of atherosclerosis. Chronic exposure to elevated lipoproteins causes accumulation of LDL in the subendothelial layer and activation of the endothelium (Libbey *et al*., 2019). This leads to an inflammatory cascade that causes a cycle of leukocyte recruitment and inflammatory activation. In this pathological context, ECs are exposed to a unique microenvironment composed of relatively high concentrations of cholesterol and cytokines. Thus, elucidating how the endothelium in vivo responds to the loss of *Srebf2* in mouse models of acute and chronic inflammation will be important delineate the role of EC cholesterol homeostasis in vascular health and disease.

## Materials and Methods

### Mammalian Cell Culture

HUVECs were obtained from the Yale School of Medicine, Vascular Biology and Therapeutics Core facility. Cells were cultured in EGM-2 media (Lonza) with 10% fetal bovine serum (FBS), penicillin/streptomycin and glutamine (2.8 mM) in a 37°C incubator with 5% CO2 supply.

### RNA Sequencing

RNA was isolated using the RNeasy Plus Kit (Qiagen) and purity of total RNA per sample was verified using the Agilent Bioanalyzer (Agilent Technologies, Santa Clara, CA). RNA sequencing was performed through the Yale Center for Genome Analysis using an Illumina HiSeq 2000 platform (paired-end 150bp read length). Briefly, rRNA was depleted from RNA using Ribo-Zero rRNA Removal Kit (Illumina). RNA libraries were generated from control cells using TrueSeq Small RNA Library preparation (Illumina) and sequenced for 45 cycles on Illumina HiSeq 2000 platform (paired end, 150bp read length).

### RNA-seq Analysis

Normalized counts and gene set enrichment analysis statistics were generated with Partek Flow. Reads were aligned to the hg19 build of the human genome with STAR and quantified to an hg19 RefSeq annotation model through Partek E/M. Gene counts were normalized as counts per million (CPM) and differential analysis was performed with GSA. Ingenuity Pathway Analysis (Ingenuity Systems QIAGEN) software was used to perform Canonical Pathway and Upstream Regulator analyses (Cutoff: p<0.05; −1.5>Fold Change>1.5). Metabolic network analysis was done using MetaCore (Clarivate) (Cutoff: p<0.005). GSEA analysis was used to produce Hallmark gene sets (1000 permutations, collapse to gene symbols, permutate to phenotype). Data are deposited in NCBI Gene Expression Omnibus and are available under GEO accession GSE201466.

### Western Blotting Analysis

Cells or tissues were lysed on ice with ice-cold lysis buffer containing 50 mM Tris-HCl, pH 7.4, 0.1 mM EDTA, 0.1 mM EGTA, 1% Nonidet P-40, 0.1% sodium deoxycholate, 0.1% SDS, 100 mM NaCl, 10 mM NaF, 1 mM sodium pyrophosphate, 1 mM sodium orthovanadate, 1 mM Pefabloc SC, and 2 mg/ml protease inhibitor mixture (Roche Diagnostics) and samples prepared. Total protein (25μg) was loaded into SDS-PAGE followed by transfer to nitrocellulose membranes. Immunoblotting was performed at 4°C overnight followed by 1hr incubation with LI-COR compatible fluorescent-labeled secondary antibodies (LI-COR Biosciences). Bands were visualized on the Odyssey CLx platform (LICOR Biosciences). Quantifications were based on densitometry using ImageJ.

### Quantitative RT-qPCR

RNA from cells or tissues were isolated using the RNeasy Plus Kit (Qiagen). 0.5 mg RNA/sample was retrotranscribed with the iScript cDNA Synthesis Kit (BioRad). Real-time quantitative PCR (qPCR) reactions were performed in duplicate using the CFX-96 Real Time PCR system (Bio-Rad). Quantitative PCR primers were designed using Primer3 software and synthesized by Yale School of Medicine Oligo Synthesis facility. Fold changes were calculated using the comparative Ct method.

### DiI-LDL Uptake

Cells were washed in PBS and treated for 1hr with plain EBM-2 containing 2.5μg/mL DiI-LDL (Kalen Biomedical). Cells were washed for 5min with acid wash (25mM Glycine, 3% (m/V) BSA in PBS at pH 4.0), before suspended in PBS, washed, and fixed. PE mean fluorescence intensity per cell was measured by LSRII (BD Biosciences) flow cytometer the same day of the assay and analyzed using FlowJo.

### Thin Layer Chromatography (TLC)

Dried lipids were resuspended in hexane and loaded onto a silica gel TLC 60 plate (Millipore Sigma) and run in hexane:diethyl ether:acetic acid (70:30:1) until the solvent line reached approximately 1 inch from the top. Standards of pure triglycerides, diacylglycerides, cholesterol, and cholesterol ester were loaded for reference. After drying, the plate was exposed to a phosphor screen for 1 week and imaged using a Typhoon phosphorimager.

### Cholesterol Efflux Assay

Cells were equilibrated with 1μCi/mL 3H-cholesterol (PerkinElmer) for 16hrs in full media containing FBS and ACAT inhibitor 58035 (Sigma). Next, cells were washed twice with PBS and incubated for 6hrs in serum-free media containing 58035 and indicated cholesterol acceptor. Media and cell lysis were harvested at the end of 6 hr. Ultima Gold scintillation liquid (PerkinElmer) were added to the media and cell lysis, respectively, and radioactivity was quantified using a Tri-Carb 2100 liquid scintillation counter (PerkinElmer). Efflux was measured as percent counts in media divided by counts in the cell lysis.

### Filipin Staining

Confluent HUVEC cells were fixed and stained with 50μg/mL Filipin and FITC-conjugated lectin from Ulex Europaeus Agglutinin I (FITC-UEAI). Images were taken on a confocal microscope (SP5, Leica). UV signal (Filipin) was immediately recorded after FITC-UEAI was used to find appropriate z-stack/cellular context.

### Total Cholesterol Extraction and Quantification

Total lipids extracted in 2:1 chloroform methanol. The solution was dried under nitrogen gas. Cholesterol was quantified according to the kit protocol (abcam).

#### Lipidomics

Mass spectrometry-based lipid analysis was performed by Lipotype GmbH (Dresden, Germany) as described (Sampaio et al. 2011). Lipids were extracted using a two-step chloroform/methanol procedure (Ejsing et al. 2009). Samples were spiked with internal lipid standard mixture containing: cardiolipin 14:0/14:0/14:0/14:0 (CL), ceramide 18:1;2/17:0 (Cer), diacylglycerol 17:0/17:0 (DAG), hexosylceramide 18:1;2/12:0 (HexCer), lyso-phosphatidate 17:0 (LPA), lyso-phosphatidylcholine 12:0 (LPC), lyso-phosphatidylethanolamine 17:1 (LPE), lyso-phosphatidylglycerol 17:1 (LPG), lyso-phosphatidylinositol 17:1 (LPI), lyso-phosphatidylserine 17:1 (LPS), phosphatidate 17:0/17:0 (PA), phosphatidylcholine 17:0/17:0 (PC), phosphatidylethanolamine 17:0/17:0 (PE), phosphatidylglycerol 17:0/17:0 (PG), phosphatidylinositol 16:0/16:0 (PI), phosphatidylserine 17:0/17:0 (PS), cholesterol ester 20:0 (CE), sphingomyelin 18:1;2/12:0;0 (SM), sulfatide d18:1;2/12:0;0 (Sulf), triacylglycerol 17:0/17:0/17:0 (TAG) and cholesterol D6 (Chol). After extraction, the organic phase was transferred to an infusion plate and dried in a speed vacuum concentrator. 1st step dry extract was re-suspended in 7.5 mM ammonium acetate in chloroform/methanol/propanol (1:2:4, V:V:V) and 2nd step dry extract in 33% ethanol solution of methylamine in chloroform/methanol (0.003:5:1; V:V:V). All liquid handling steps were performed using Hamilton Robotics STARlet robotic platform with the Anti Droplet Control feature for organic solvents pipetting. Samples were analyzed by direct infusion on a QExactive mass spectrometer (Thermo Scientific) equipped with a TriVersa NanoMate ion source (Advion Biosciences). Samples were analyzed in both positive and negative ion modes with a resolution of Rm/z=200=280000 for MS and Rm/z=200=17500 for MSMS experiments, in a single acquisition. MSMS was triggered by an inclusion list encompassing corresponding MS mass ranges scanned in 1 Da increments (Surma et al. 2015). Both MS and MSMS data were combined to monitor CE, DAG and TAG ions as ammonium adducts; PC, PC O-, as acetate adducts; and CL, PA, PE, PE O-, PG, PI and PS as deprotonated anions. MS only was used to monitor LPA, LPE, LPE O-, LPI and LPS as deprotonated anions; Cer, HexCer, SM, LPC and LPC O-as acetate adducts and cholesterol as ammonium adduct of an acetylated derivative (Liebisch et al. 2006). Data were analyzed with in-house developed lipid identification software based on LipidXplorer (Herzog et al. 2011; Herzog et al. 2012). Data post-processing and normalization were performed using an in-house developed data management system. Only lipid identifications with a signal-to-noise ratio >5, and a signal intensity 5-fold higher than in corresponding blank samples were considered for further data analysis.

#### ALOD4 and OlyA Purification

ALOD4 and OlyA expression constructs were generously provided by the lab of Dr. Arun Radhakrishnan. Recombinant His-tagged ALOD4 and OlyA were purified as previously described (Endapally *et al.,* 2019). Briefly, ALOD4 expression was induced with 1mM IPTG in OD0.5 BL21 (DE3) pLysS *E. coli* for 16hr at 18°C. Cells were lysed and His-ALOD4 and His-OlyA were isolated by nickel purification followed by size exclusion chromatography (HisTrap-HP Ni column, Tricorn 10/300 Superdex 200 gel filtration column; FPLC AKTA, GE Healthcare). Protein-rich fractions were pooled and concentration was measured using a NanoDrop instrument.

#### ALOD4 Fluorescent Labeling

20nmol ALOD4 was combined with 200nm AlexaFluor maleimide (ThermoFisher) in 50mM Tris-HCl, 1mM TCEP, 150mM NaCl pH 7.5 and incubated at 4°C for 16hr. The reaction was quenched using 10mM DTT. Unbound fluorescent label and DTT were removed by dialysis (EMD Millipore).

#### ALOD4 Binding and Western Blot Analysis

At time of collection, HUVEC were washed 3 times for 5min in PBS with Ca^2+^ and Mg^2+^ containing 0.2% (wt/vol) BSA. Cells were then incubated with 3μM ALOD4 in basal EBM2 media containing 0.2% (wt/vol) BSA for 1hr at 4°C. The unbound proteins were removed by washing three times with PBS with Ca^2+^ and Mg^2+^ for 5min each. Cells were then lysed and prepared for SDS-PAGE and immunoblotting. ALOD4 was probed on nitrocellulose gels using anti-6X His (abcam) antibody at 15kDa. A similar method was used for OlyA binding.

#### ALOD4 In-Cell Western Blot Analysis

Cells were cultured onto 96 wells and ALOD4 binding was performed as mentioned above up until lysis. Cells were directly incubated with DyLight680-conjugated anti-His antibody (Thermofisher), washed, and 700nm fluorescence was recorded directly on Odyssey CLx platform (LICOR Biosciences).

#### ALOD4 Flow Cytometry Analysis

Cells were suspended in PBS with Ca^2+^ and Mg^2+^ containing 2% FBS and washed 3 times. Binding was with 3μM ALOD4-647 for 1hr at 4°C. Cells were then washed 3 times with PBS with Ca^2+^ and Mg^2+^ containing 2% FBS and mean fluorescence intensity per cell was measured by LSRII (BD Biosciences) flow cytometer the same day of the assay.

#### Animal Studies

All animals were handed according to approved institutional animal care and use committee (IACUC) protocols (#07919-2020) of Yale University. At 10 weeks of age, male C57BL/6J mice (JAX, #000664) were injected with 15mg/kg lipopolysaccharide (LPS) from E. Coli O111:B4 intraperitoneally (Sigma). 6 hours later, blood was collected for lipid and cytokine analysis. Mice were perfused with PBS and lungs were processed for flow cytometry analysis. Briefly, lung cells were brought to a single-cell suspension via collagenase incubation and then stained for flow cytometry at a concentration of 5×10^6^ cells/mL with Cd31 (Biolegend) and 3μM ALOD4-47.

#### Statistics

Statistical differences were measured with an unpaired 2-sided Student’s t-test or ANOVA with listed correction for multiple corrections. A value of p<0.05 was considered statistically significant. “n” within figure legends involving HUVEC denotes number of donors used for the respective experiment. Data analysis was performed with GraphPad Prism software (GraphPad, San Diego, CA).

#### Oligonucleotides

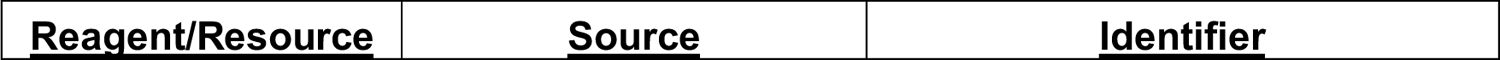
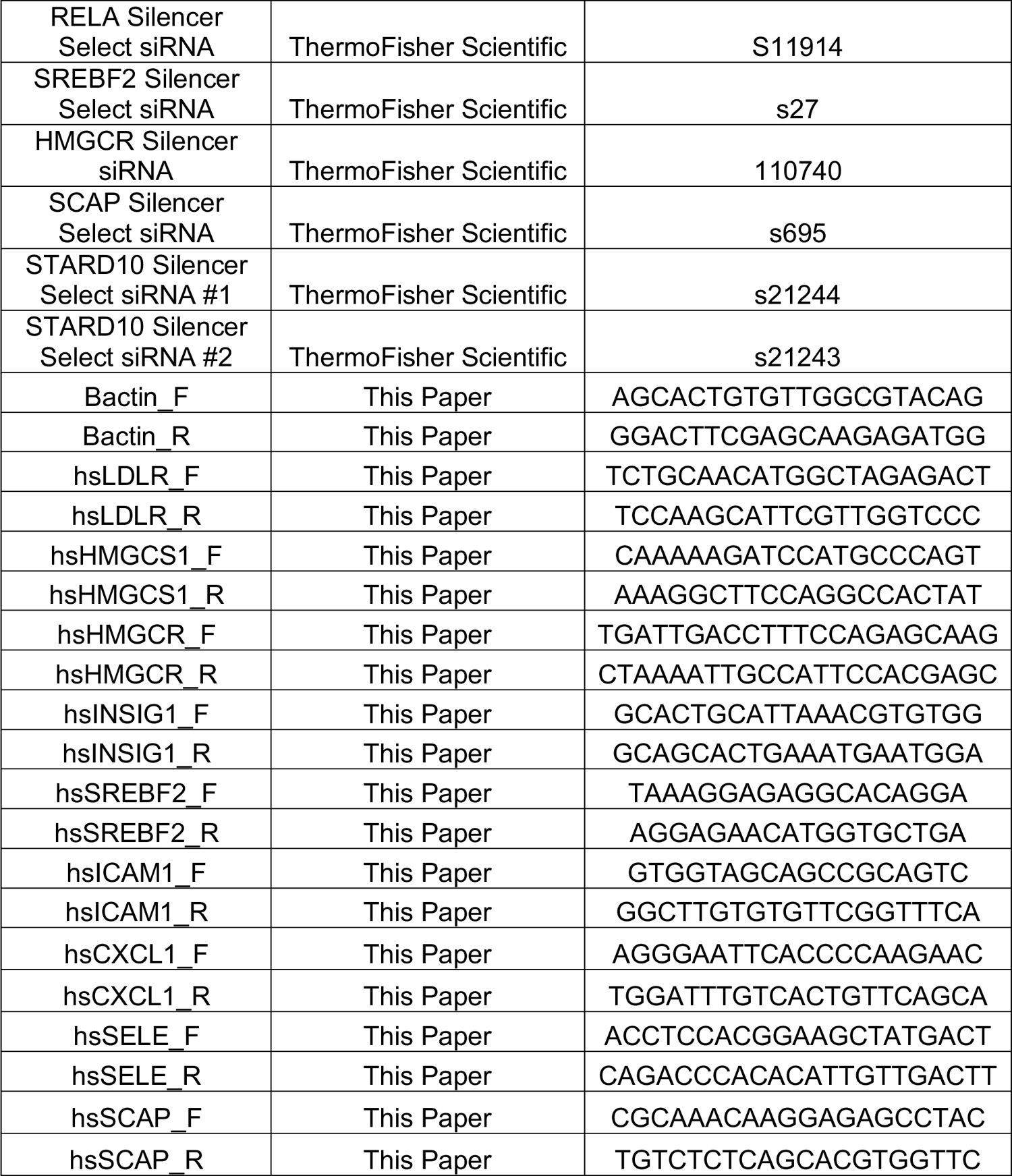

#### Antibodies

| **<u>Reagent/Resource</u>** | **<u>Source</u>** | **<u>Identifier</u>** |
| --- | --- | --- |
| 6x His Tag DyLight<br>680 | ThermoFisher<br>Scientific | MA1-21315-D680 |
| Anti-6X His | Abcam | ab18184 |
| BV605-mCD31 | BioLegend | 102427 |
| GAPDH | Cell Signaling | 2118S |
| HSP90 | Santa Cruz | sc-13119 |
| ICAM1 | Cell Signaling | 4915S |
| JNK1 | Cell Signaling | 3708S |
| LC3b | Cell Signaling | 2775S |
| LDLR | Abcam | ab52818 |
| p-JNK1 | Cell Signaling | 9261S |
| p-p38 | Cell Signaling | 9211S |
| p38 | Cell Signaling | 9212S |
| P65 (RELA) | Cell Signaling | 8242S |
| SREBP1a | Santa Cruz | sc-13551 |
| SREBP2 | BD Biosciences | 557037 |
| VCAM1 | Santa Cruz | sc-13160 |

#### Chemicals

| **<u>Reagent/Resource</u>** | **<u>Source</u>** | **<u>Identifier</u>** |
| --- | --- | --- |
| 25-Hydroxycholesterol | Sigma Aldrich | H1015 |
| Actinomycin D | ThermoFisher | 11805017 |
| BAY 117082 | Sigma Aldrich | B556-10MG |
| Cholesterol, 1,2-3H(N) | Perkin Elmer | NET139250UC |
| Choroquine (CQ) | Sigma Aldrich | C6628 |
| Dil LDL | Kalen Biomedical | 770230 |
| EGM2 | Lonza | CC-3162 |
| Fatostatin | Cayman | 13562 |
| Filipin | Cayman | 70440 |
| FITC-UEAI | ThermoFisher | L32476 |
| Lipopolysaccharide from E. coli O111:B4 | Sigma Aldrich | L2630 |
| Lipoprotein Depleted Serum (LPDS) | Kalen Biomedical | 880100 |
| MβCD | Sigma Aldrich | C4555 |
| MβCD-Cholesterol | Sigma Aldrich | C4951 |
| Native LDL | Kalen Biomedical | 770200 |
| Pefabloc | Sigma Aldrich | 11873601001 |
| PF-429242 | Sigma Aldrich | SML0667 |
| Protease Inhibitor Mixture | Roche | 11697498001 |
| rhIL1β | RD Systems | 201-LB-010/CF |
| rhTNFα | RD Systems | 210-TA-020/CF |
| Sandoz 58-035 | Sigma Aldrich | S9318-25mg |
| TLC Silica Gel 60 | Millipore Sigma | 105553 |
| Sphingomyelinase | Sigma Aldrich | S8633 |
| T0901317 | Sigma Aldrich | T2320 |
| Triacin C | RD Systems | 2472 |
| U18666A | Sigma Aldrich | U3633 |

## Acknowledgements

This work was supported by NIH grant R35HL139945, RO1DK125492, PO1 HL1070205 to WCS and a Supplement to R35HL139945 to JWF and K01DK124441 to NEB.

## Author Contributions

JWF and WCS conceived the project, designed experiments, and wrote the manuscript. RZ produced and purified ALOD4 and OlyA Probes. NEB and BT contributed intellectually and experimentally to the MS. WCS supervised the project.

## Supplemental Figure Legend

**Figure 1 – Figure Supplement 1.**
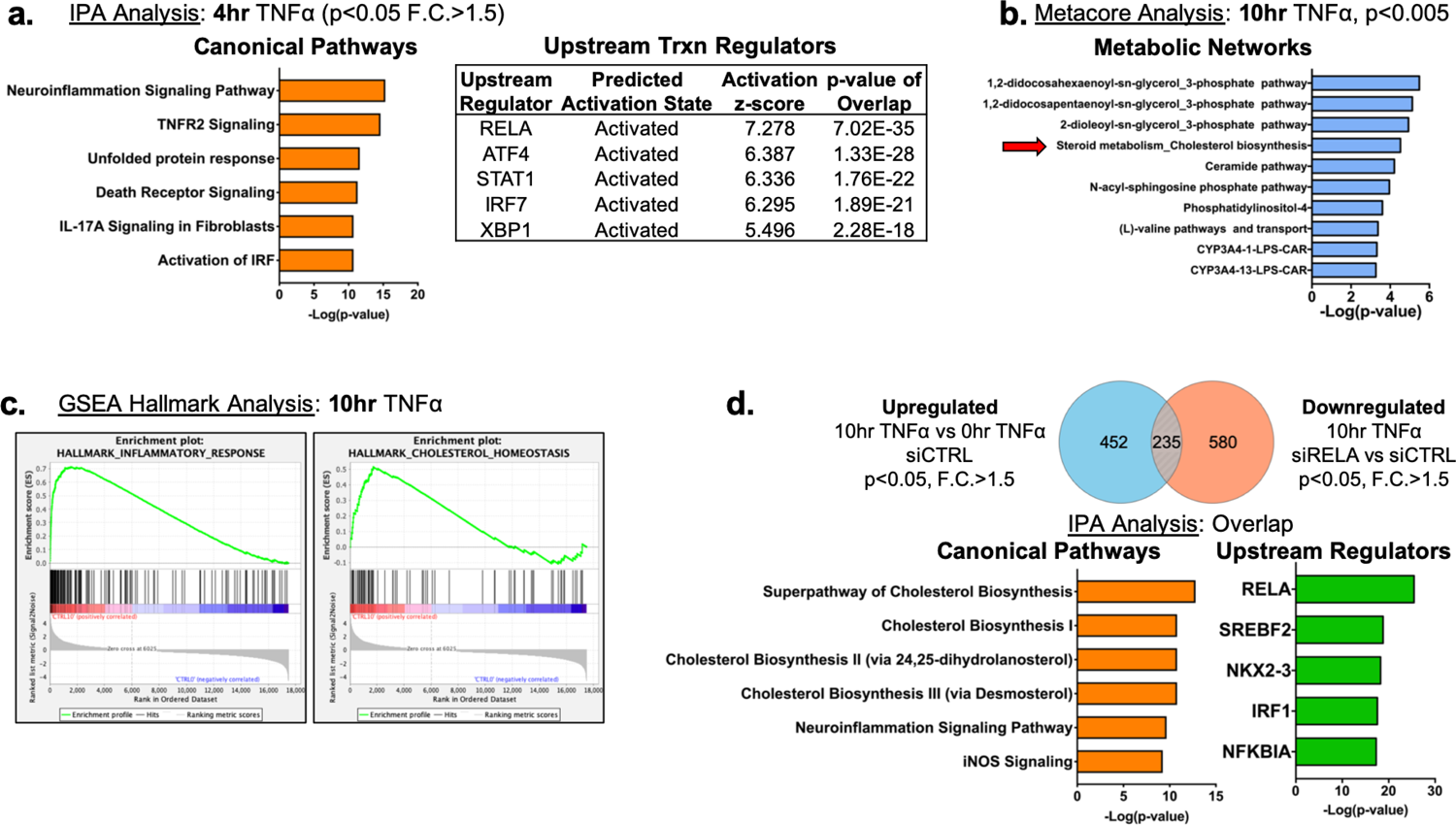
(a) Ingenuity pathway analysis for pathways and upstream transcription regulators using differentially expressed genes (upregulated) in HUVEC after 4 hr TNF*α* treatment (F.C.>1.5; p<0.05) (b) Metacore metabolic network analysis using upregulated genes from (Fig. 1a) (p<0.005). (c) GSEA hallmark analysis using upregulated genes from (Fig. 1a). (d) Ingenuity pathway analysis of gene set overlap between significantly upregulated genes in 10 hr TNF*α* compared to 0 hr TNF*α* and significantly downregulated genes after 10hr TNF*α* and in siRELA compared to siCTRL.

**Figure 2 – Figure Supplement 1.**
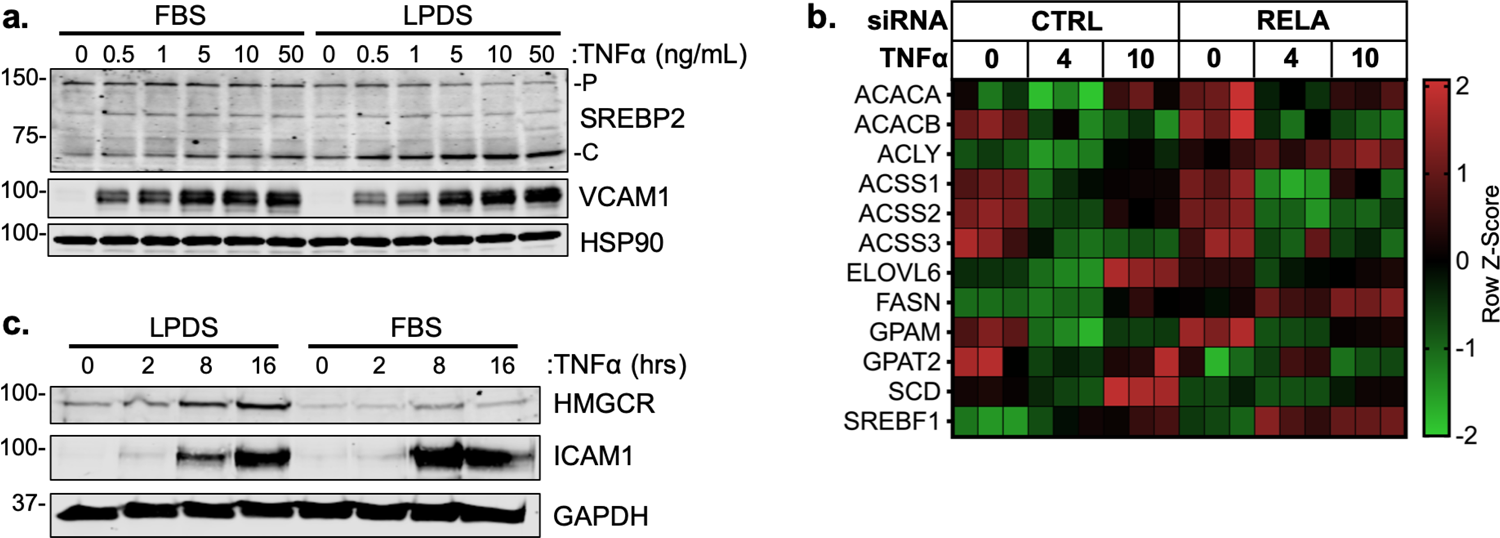
(a) Representative SREBP2 immunoblot from whole-cell lysates from HUVEC treated with TNF*α* (16 hr) at indicated dose. Cells were incubated with fetal bovine serum (FBS) or lipoprotein depleted serum (LPDS). (b) Heatmap of classical SREBP1-dependent fatty acid synthesis genes from previous RNA-seq analysis. (c) Representative HMGCR immunoblot of HUVEC treated with TNF*α*(10ng/mL) for indicated time and media.

**Figure 4 – Figure Supplement 1.**
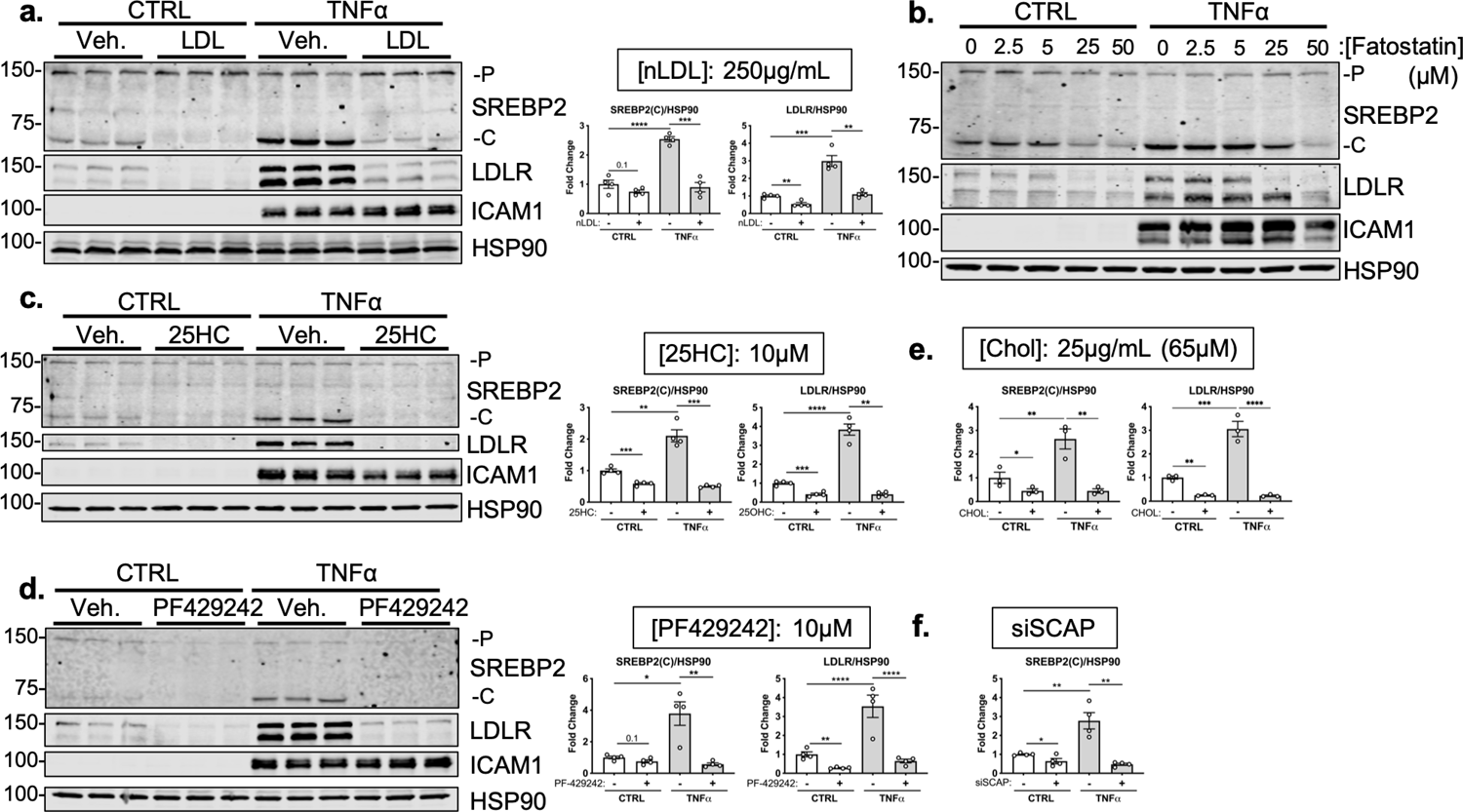
(a) SREBP2 and LDLR protein levels in HUVEC treated with TNF*α* (10ng/mL) and with or without low density lipoprotein (LDL) (250μg/mL). Data are normalized to respective HSP90 and then to untreated cells (n=4). (b) Representative immunoblot of SREBP2 and LDLR protein levels in HUVEC treated with TNF*α* (10ng/mL) and increasing concentrations of fatostatin. (c) SREBP2 and LDLR protein levels in HUVEC treated with TNF*α* (10ng/mL) and with or without 25-hydroxycholesterol (25HC) (10μM). Data are normalized to respective HSP90 and then to untreated cells (n=4). (d) SREBP2 and LDLR protein levels in HUVEC treated with TNF*α* (10ng/mL) and with or without PF-429242 (10μM). Data are normalized to respective HSP90 and then to untreated cells (n=4). (e) Quantification of SREBP2 and LDLR protein levels in HUVEC treated with TNF*α* (10ng/mL) and with or without M*β*CD-cholesterol (Chol) (65μM). Data are normalized to respective HSP90 and then to untreated cells (n=3). (f) Quantification of SREBP2 protein levels in HUVEC treated with TNF*α* (10ng/mL) and with or without siSCAP. Data are normalized to respective HSP90 and then to untreated cells (n=4). *p<0.05; **p<0.01; ***p<0.001; ***p<0.0001 by two-way ANOVA with Sidak’s multiple comparisons test

**Figure 5 – Figure Supplement 1.**
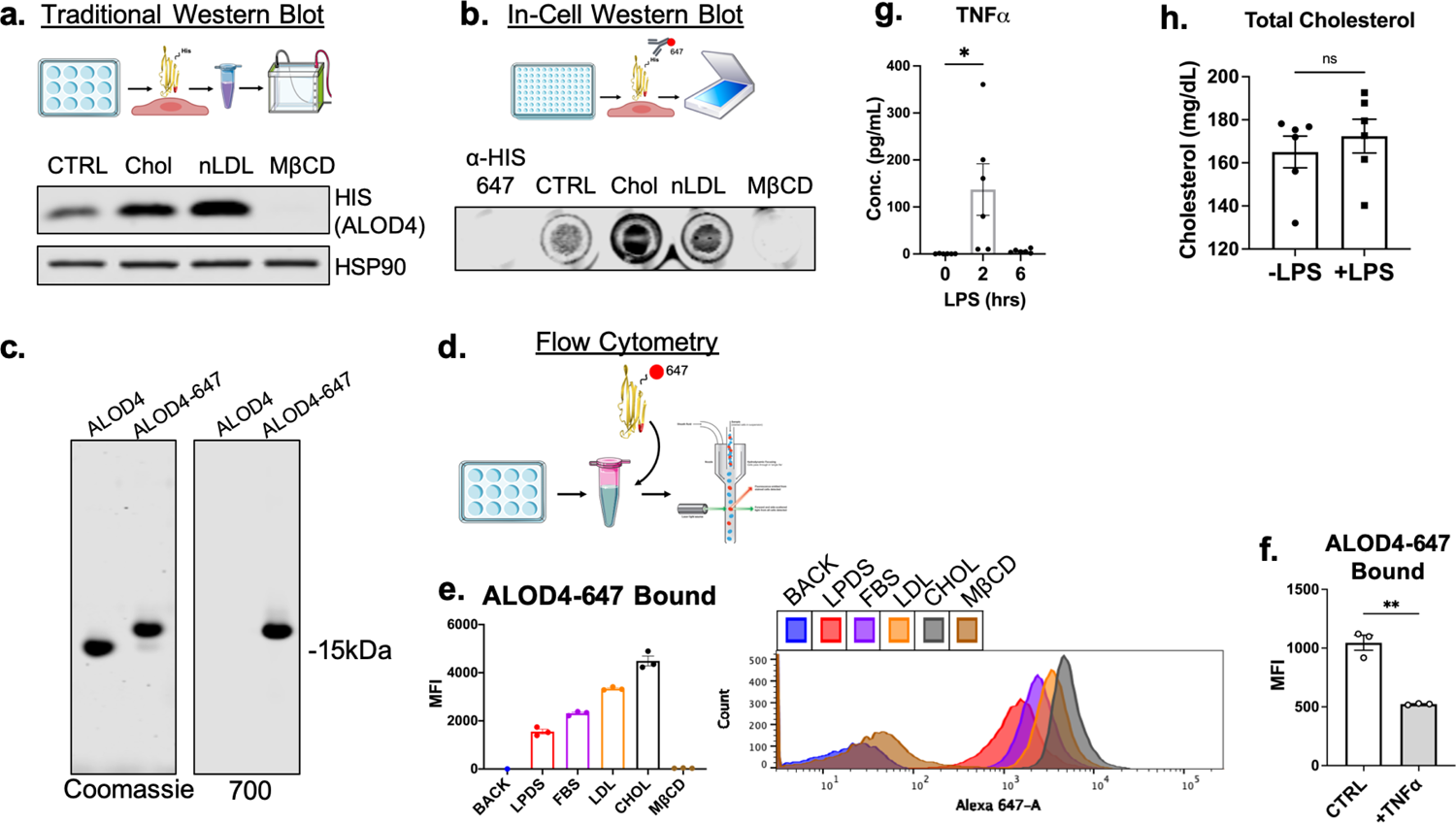
(a) Diagram of pipeline for immunoblotting protocol to quantify EC accessible cholesterol (top). Representative immunoblot of HIS (ALOD4) after treatment with cholesterol modifying agents: M*β*CD-cholesterol (Chol) (25μg/mL), LDL (100μg/ml), or M*β*CD (1%) (bottom). (b) Diagram of pipeline for in-cell Western blotting protocol to quantify EC accessible cholesterol (top). Representative in-cell Western blot of secondary alone (*α*-HIS-647) or HIS (ALOD4) after treatment with cholesterol modifying agents: M*β*CD-cholesterol (Chol) (25μg/mL), LDL (100μg/ml), or M*β*CD (1%) (bottom). (c) Representative SDS-PAGE gel of purified unconjugated ALOD4 and fluorescent ALOD4-647 stained with Coomassie (left) or recorded with the 700nm channel on LICOR Biosciences Odyssey CLx platform. (d) Schematic of flow cytometry pipeline to quantify ALOD4 binding in cultured ECs with ALOD4-647. (e) Flow cytometry analysis of bound ALOD4-647 per HUVEC after treatment with positive controls, lipoprotein depleted serum (LPDS), fetal bovine serum (FBS), LDL (100μg/mL), M*β*CD-cholesterol (Chol) (25μg/mL), or M*β*CD (1%). ALOD4 binding was quantified by mean fluorescence intensity of AlexaFluor647 channel (10,000 events/replicate, n=3). (f) Flow cytometry analysis of ALOD4-647 bound to HUVEC treated with TNF*α* (10ng/mL) for 16 hr. ALOD4 binding was quantified by mean fluorescence intensity of AlexaFluor647 channel (10,000 events/replicate, n=3). (g) Circulating TNF*α* from serum of mice treated with LPS (15mg/kg) for 2 or 6 hr (n=6). (h) Total cholesterol from serum of mice used in (Fig 5g) (n=6). *p<0.05; **p<0.01; ***p<0.001; ***p<0.0001 by unpaired t-test (f and g) or one-way ANOVA with Tukey’s multiple comparison’s test.

**Figure 6 – Figure Supplement 1.**
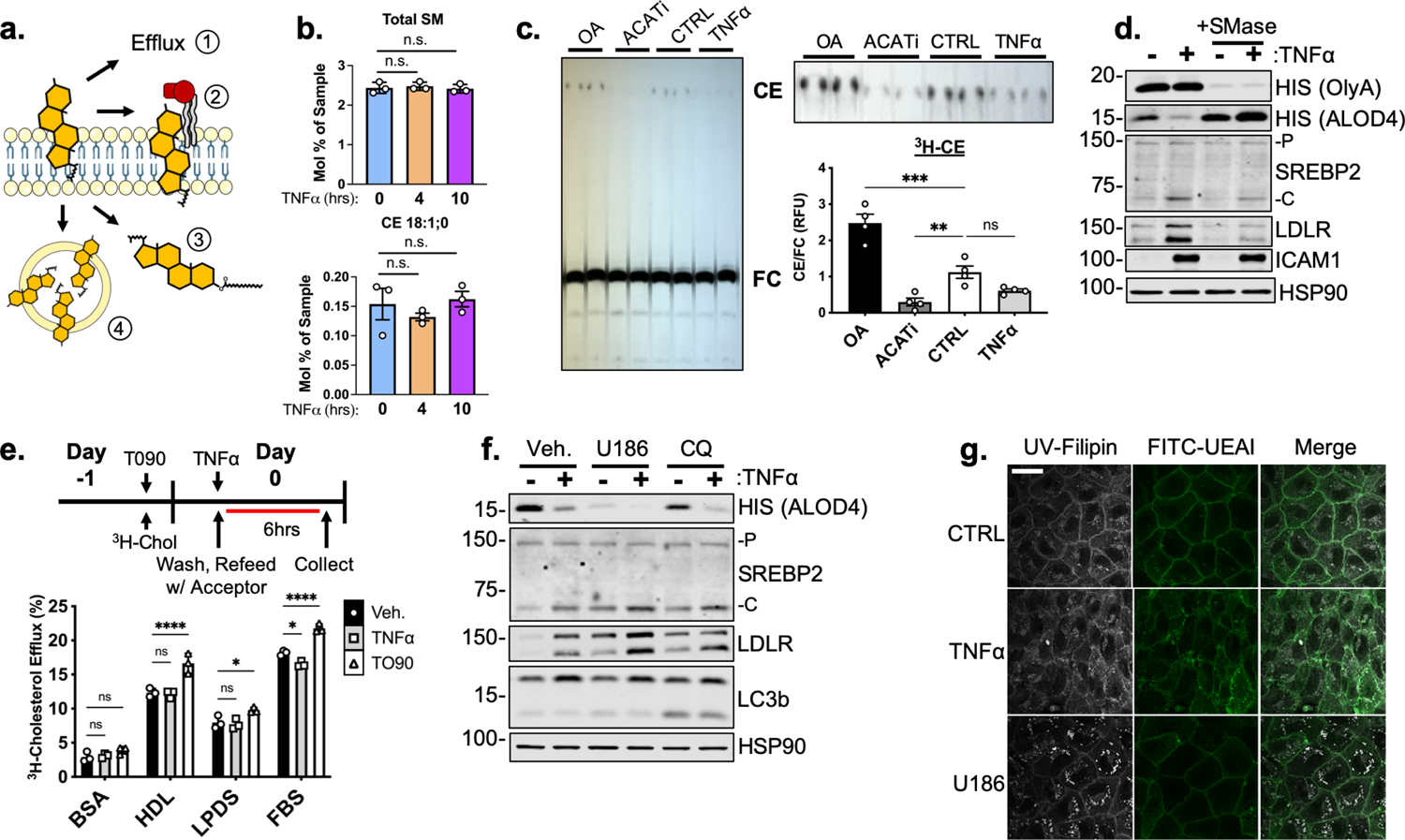
(a) Schematic of possible mechanisms to deplete plasma membrane accessible cholesterol: (1) efflux, (2) sphingomyelin shielding, (3) esterification, and (4) lysosomal/endosomal accumulation. (b) Total sphingomyelin (SM) and cholesteryl ester (CE) content in HUVEC after 4 or 10 hr of TNF*α* (10ng/mL) quantified by mass spectrometry (n=3). (c) Thin layer chromatography of 3H-cholesterol isolated from HUVEC treated with oleic acid (OA) (0.5mM), Sandoz 58-035 (ACATi) (1μM), or TNF*α* (10ng/mL) for 16hr. Esterification was quantified as a ratio between cholesteryl ester (CE) and free cholesterol (FC) (n=4). (d) Representative immunoblot of OlyA, ALOD4, SREBP2, and LDLR protein levels in HUVEC treated with TNF*α* (10ng/mL) and sphingomyelinase (SMase) (100mU/mL). (e) Schematic of protocol for measurement of cholesterol efflux (top). 3H-cholesterol efflux in HUVEC treated with T0901317 (T090) (5μM) or TNF*α* (10ng/mL) and with indicated acceptors, BSA, HDL, lipoprotein depleted serum (LPDS), or fetal bovine serum (FBS). Efflux was quantified as the ratio of 3H-cholesterol in the media compared to lysates (n=4). (f) Immunoblot of ALOD4, SREBP2, and LDLR protein levels in HUVEC treated with U18666A (U186) (5μM) or choloroquine (CQ) (10μM) and with or without TNF*α* (10ng/mL). Data are normalized to respective HSP90 and then to untreated cells (n=3). (g) Representative images of Filipin and FITC-ulex eruopaeus agglutinin I (UEAI) stained HUVEC after treatment with TNF*α* (10ng/mL) or U18666A (U186) (5μM). White scale bar = 30 μm. *p<0.05; **p<0.01; ***p<0.001; ***p<0.0001 by one-way ANOVA with Tukey’s multiple comparison’s test (c and e)

## Supplemental Files Legends

**Supplementary File 1. RNA-seq normalized counts and lipidomics** (RNAseq) HUVEC were treated with TNF*α*for 0, 4, and 10 hr and with or without siRNA targeting *RELA*. (Lipidomics). HUVEC were treated with TNF*α*for 0, 4, and 10 hr. Data represented as molar percentage of lipid

## Source Data Legends

**Figure 2_Source Data 1.** Blots corresponding to Figure 2a and 2c.

**Figure 2_Source Data 2.** Raw data supporting Figure 2a, 2b, 2c, and 2d

**Figure 2 – Figure Supplement 1_Source Data 1.** Blots corresponding to Figure 2 – Figure Supplement 1 a and c.

**Figure 3_Source Data 1.** Blots corresponding to Figure 3a, 3b, 3c, and 3e.

**Figure 3_Source Data 2.** Raw data supporting Figure 3c, 3d, and 3e.

**Figure 4_Source Data 1.** Blots corresponding to Figure 4b, 4c, 4d, 4e, and 4f.

**Figure 4_Source Data 2.** Raw data supporting Figure 4g and Figure 4 – Supplement 1 a, c, d, e, and f.

**Figure 4 – Figure Supplement 1_Source Data 1.** Blots corresponding to Figure 4 – Figure Supplement 1 a, b, c, and d.

**Figure 5_Source Data 1.** Blots corresponding to Figure 5c and 5f.

**Figure 5_Source Data 2.** Raw data supporting Figure 5a, 5c, 5d, 5e, 5f, 5i and Figure 5 – Figure Supplement 1 e, f, g, and h.

**Figure 5 – Figure Supplement 1_Source Data 1.** Blots corresponding to Figure 5 – Figure Supplement 1 a and c.

**Figure 6_Source Data 1.** Blots corresponding to Figure 6e.

**Figure 6_Source Data 2.** Raw data supporting Figure 6d and 6e and Figure 6 – Figure Supplement 1 c and e.

**Figure 6 – Figure Supplement 1_Source Data 1.** Blots corresponding to Figure 6 – Figure Supplement 1 c, d, and f.

